# Metabolic differences between symbiont subpopulations in the deep-sea tubeworm *Riftia pachyptila*

**DOI:** 10.1101/2020.04.08.032177

**Authors:** Tjorven Hinzke, Manuel Kleiner, Mareike Meister, Rabea Schlüter, Christian Hentschker, Jan Pané-Farré, Petra Hildebrandt, Horst Felbeck, Stefan M. Sievert, Florian Bonn, Uwe Völker, Dörte Becher, Thomas Schweder, Stephanie Markert

## Abstract

The hydrothermal vent tube worm *Riftia pachyptila* lives in intimate symbiosis with intracellular sulfur-oxidizing gammaproteobacteria. Although the symbiont population consists of a single 16S rRNA phylotype, bacteria in the same host animal exhibit a remarkable degree of metabolic diversity: They simultaneously utilize two carbon fixation pathways and various energy sources and electron acceptors. Whether these multiple metabolic routes are employed in the same symbiont cells, or rather in distinct symbiont subpopulations, was unclear. As *Riftia* symbionts vary considerably in cell size and shape, we enriched individual symbiont cell sizes by density gradient centrifugation in order to test whether symbiont cells of different sizes show different metabolic profiles. Metaproteomic analysis and statistical evaluation using clustering and random forests, supported by microscopy and flow cytometry, strongly suggest that *Riftia* symbiont cells of different sizes represent metabolically dissimilar stages of a physiological differentiation process: Small symbionts actively divide and may establish cellular symbiont-host interaction, as indicated by highest abundance of the cell division key protein FtsZ and highly abundant chaperones and porins in this initial phase. Large symbionts, on the other hand, apparently do not divide, but still replicate DNA, leading to DNA endoreduplication. Highest abundance of enzymes for CO_2_ fixation, carbon storage and biosynthesis in large symbionts indicates that in this late differentiation stage the symbiont’s metabolism is efficiently geared towards the production of organic material. We propose that this division of labor between smaller and larger symbionts benefits the productivity of the symbiosis as a whole.

## Introduction

The chemoautotrophic gammaproteobacterium *Candidatus* Endoriftia persephone, sulfur-oxidizing endosymbiont of the deep-sea tubeworm *Riftia pachyptila*, provides all nutrition for its gutless host (Cavanaugh *et al.*, 1981, Felbeck, 1981, Hand, 1987, Distel and Felbeck, 1988, Robidart *et al.*, 2008). *Ca.* E. persephone (here Endoriftia) densely populates *Riftia*’s trophosome, a specialized organ in the worm’s trunk, where the bacteria are housed intracellularly in host bacteriocytes (Hand, 1987).

Although the symbiont population consists of a single 16S rRNA phylotype (Polzin *et al.*, 2019), it was previously shown to exhibit remarkable metabolic versatility: As demonstrated by proteomic analyses, symbionts from the same host animal expressed enzymes of two CO_2_ fixation pathways, the Calvin cycle and the reverse tricarboxylic acid (rTCA) cycle, as well as enzymes for both, glycogen generation and glycogen degradation (Markert *et al.*, 2007, Markert *et al.*, 2011, Gardebrecht *et al.*, 2012, Hinzke *et al.*, 2019). Moreover, proteins involved in utilization of hydrogen sulfide and thiosulfate as energy sources were expressed simultaneously by the same symbiont population; as were proteins for the use of nitrate and oxygen as electron acceptors (Markert *et al.*, 2011). Based on these observations, we hypothesized that individual, metabolically distinct symbiont subpopulations in the trophosome may exist.

These presumptive subpopulations are likely congruent with symbionts of different cell sizes: Individual Endoriftia cells exhibit pronounced morphological diversity, ranging from small rods to small and large cocci in ultimate proximity to each other within the same host specimen (Hand, 1987, Bright *et al.*, 2000, Bright and Sorgo, 2003). In individual trophosome lobules, which measure approximately 200-500 µm in diameter, the smallest, rod-shaped symbiont cells are located close to the central blood vessel, while towards the lobule periphery, symbionts gradually increase in size and become coccoid, before they are degraded in the outermost lobule zone. Only small Endoriftia cells and the host bacteriocytes in which they reside appear to undergo cell division, indicating that small and large symbionts belong to a common cell cycle (Bright *et al.*, 2000, Bright and Sorgo, 2003). Previous microscopy-based studies indicated that small and large *Riftia* symbionts differ not only with regard to their frequency of cell division, but also with regard to carbon incorporation rates, amount of stored glycogen, and area of sulfur storage vesicles (Bright *et al.*, 2000, Sorgo *et al.*, 2002, Bright and Sorgo, 2003, Pflugfelder *et al.*, 2005). This suggests that individual cell sizes may indeed have dissimilar metabolic properties.

In this study, we aimed to analyze and compare the metabolic profiles of individual *Riftia* symbiont subpopulations. In contrast to previous molecular analyses that studied metabolic capabilities of the *Riftia* symbiont population as a mixture of all cell sizes (e.g., Markert *et al.*, 2007, Markert *et al.*, 2011, Gardebrecht *et al.*, 2012), precluding comparisons between putative subpopulations, we used a more sensitive approach: We enriched Endoriftia cells of different sizes by gradient centrifugation of trophosome tissue homogenate and subjected these enriched gradient fractions to separate metaproteomic analyses. Statistical evaluation using clustering and random forests allowed us to deduce cell size-dependent differences in protein abundance and metabolic functions. Catalyzed reporter deposition-fluorescence *in situ* hybridization (CARD-FISH), transmission electron microscopy (TEM), hybridization chain reaction (HCR)-FISH analyses, and flow cytometry complemented these experiments. Our results suggest a division of labor between different developmental stages of the symbiont.

## Material and Methods

### Sample collection and enrichment of symbiont subpopulations

*Riftia* samples for enrichment of symbiont subpopulations were collected at the East Pacific Rise hydrothermal vent field at 9°50’ N, 104°17’ W in a water depth of about 2,500 m during a research cruise with R/V Atlantis in November 2014 (AT26-23). Samples for electron microscopy were obtained during a second cruise (AT37-12) at the same site during March-April 2017 (Hinzke *et al.*, 2019). Sample details and numbers of biological replicates are summarized in Supplementary Table S1.

Trophosome sulfur content of the specimens was estimated based on the trophosome tissue’s color: sulfur-rich (S-rich) specimens have a light yellowish trophosome, due to the sulfur stored in the symbionts, whereas trophosomes of sulfur-depleted (S-depleted) specimens appear dark green to black (Pflugfelder *et al.*, 2005). For proteomic analyses, we used four S-rich *Riftia* specimens and three S-depleted specimens.

To enrich symbiont cells of varying sizes (i.e., morphologically distinct symbiont subpopulations), *Riftia* specimens were dissected onboard the research vessel immediately after recovery of the worms and approximately 3 ml trophosome tissue were homogenized in a Dounce glass homogenizer in 6 ml imidazole-buffered saline (IBS, 0.49 M NaCl, 0.03 M MgSO^4^, 0.011 M CaCl2, 0.003 M KCl, 0.05 M imidazole). As described in Hinzke *et al.* (2018), the homogenate was subjected to rate-zonal density gradient centrifugation, which allows to separate particles based on their size (Graham, 2001). In brief, an 8-18% Histodenz™ density gradient was created using a dilution series of Histodenz™ in IBS (1% steps, 1 ml per step), which was stacked in a 15 ml tube so that Histodenz™ concentration was highest at the bottom and lowest at the top. 0.5 ml tissue homogenate was layered on top of this gradient and the gradient was centrifuged (1,000 x g, 5 min, 4 °C) in a swing-out rotor. Smaller symbiont cells were thus enriched in less dense gradient fractions (lower Histodenz™ concentrations) in the upper part of the gradient, while larger cells migrated to lower gradient fractions. After centrifugation, gradients were disassembled by carefully fractionizing the entire gradient volume into 0.5 ml subsamples, giving a total of 24 fractions. Enrichment of distinct symbiont subpopulations in these subsamples was confirmed using catalyzed reporter deposition-fluorescence in situ hybridization (CARD-FISH, see below). For this purpose, 20 µl of each gradient fraction subsample and 15 µl of homogenate was fixed in 1% PFA in IBS, and symbiont cells were subsequently filtered onto GTTP polycarbonate filters (pore size 0.2 μm, Millipore) as described previously (Ponnudurai *et al.*, 2017).

### CARD-FISH

Enrichment of symbiont cell sizes in gradient fractions was analyzed employing fluorescence microscopy with samples labelled by CARD-FISH. CARD-FISH labeling was performed as previously described (Ponnudurai *et al.*, 2017), using the probe Rif445 (Nussbaumer *et al.*, 2006) and Alexa Fluor^®^ 594-labeled tyramide. For counterstaining, 0.1% (w/v) 4,6-diamidino-2-phenylindole (DAPI) was added to the embedding medium (4:1 Citifluor AF1 (Citifluor) and Vectashield (Vector Laboratories)). CARD-FISH filters were analyzed using an Axio Imager.M2 fluorescence microscope (Carl Zeiss Microscopy GmbH). For semi-automated cell counting and to measure the longest cell dimension, we used a custom Fiji (Schindelin *et al.*, 2012) macro with the Fiji plugins Enhanced Local Contrast (CLAHE; Saalfeld, 2010) and Bi-exponential edge preserving smoother (BEEPS; Thévenaz *et al.*, 2012). After image processing, we excluded objects with a size of less than 2 µm (as these were mainly artifacts) and set the maximum object size to 20 µm. To assign cell sizes to size classes (i.e., cell size ranges) we used a quartile split: We calculated quartiles of cell sizes in non-enriched homogenate samples (i.e., 25% of all cells in homogenate samples were assigned to each class). This resulted in the four calculative size classes very small (≥2 µm – <3.912 µm), small (≥3.912 µm – <5.314 µm), medium (≥5.314 µm – <6.83275 µm) and large (≥6.83275 µm – 20 µm). The majority of cells in all size classes were coccoid. Rod-shaped cells were almost exclusively present in the smallest size class. Individual gradient fractions (subsamples) were screened for their respective share of cells in each size class and the subsample with the highest percentage of cells in the respective quartile was chosen for metaproteomic analysis. For example, if of all 24 subsamples of a sample, fraction 5 had the highest percentage of very small cells, i.e. most of the cells in fraction 5 were between 2 µm and 3.912 µm in diameter (as measured by our Fiji macro), this fraction was chosen as representative of very small symbiont cells in the respective biological replicate (worm). The fraction containing the highest percentage of very small cells will be referred to as “fraction XS” in the following. The fractions containing the highest percentages of small, medium and large symbiont cells will be referred to as “S”, “M”, and “L”, respectively. If the same subsample had the highest percentage of cells in two size classes, this subsample was chosen as representative for one of these size classes, and for the other size class, the subsample with the second highest percentage of cells in that class was used as representative. Cell size distributions in the four size class representatives are summarized in Figure 1.

**Figure 1:**
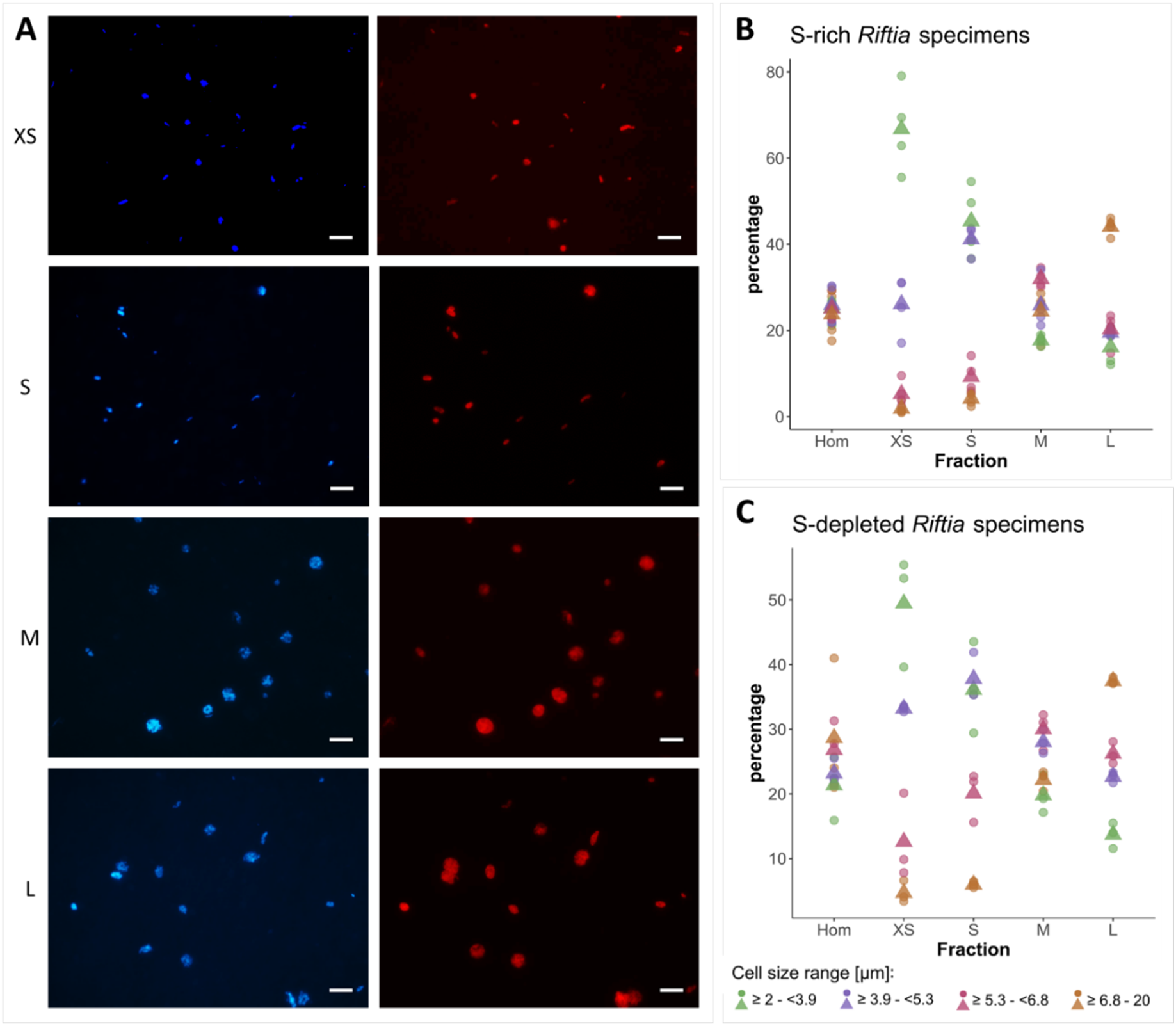
**A)** Catalyzed reporter deposition-fluorescence *in situ* hybridization (CARD-FISH) images of *Riftia* symbiont cells after density gradient centrifugation of trophosome homogenate. After the enrichment procedure, small bacterial cells had accumulated in the upper, less dense gradient fractions (top), while larger symbionts were enriched in the lower, denser fractions (bottom). Left: DAPI staining, right: 16S rRNA signal. For better visibility, brightness and contrast were adjusted in all images. **B and C)** Symbiont cell size distributions in individual gradient fractions. While all cell size groups were roughly equally abundant in non-enriched trophosome homogenate (Hom), fraction XS had the highest percentage of symbiont cells in the size range 2.0 µm - 3.9 µm, fraction S contained most symbiont cells of 3.9 µm – 5.3 µm, etc. Gradient centrifugation was performed using four biological replicates (n=4) of sulfur-rich trophosome tissue (B) and three biological replicates (n=3) of sulfur-depleted trophosome tissue (C). For an overview of which gradient fractions were chosen as fractions XS, S, M, and L in all samples see Supplementary Table S1. Dots: individual % values, triangles: average % values.

### Transmission electron microscopy (TEM)

Trophosome samples used for TEM in this study (see Supplementary Table S1 for details) were prepared and analyzed as described previously (Hinzke *et al.*, 2019). Tissue sections were recorded on sheet films (Kodak electron image film SO-163, Plano GmbH, Wetzlar) as described by Petersen *et al*. (2020). To create a composite high-resolution TEM image of a trophosome lobule (Figure 5A), we merged 50 individual micrographs of one section using Serif Affinity Photo (https://affinity.serif.com/en-us/photo/). All 50 partially overlapping images were loaded and the fully automated “Panorama Stitching” technique was applied, resulting in a panorama image still showing some vignette marks caused by inhomogeneous exposure at the former edges of individual images. The global smooth frequencies reflecting these exposure errors were removed using the frequency separation filter with a large radius. The gradation curve was manually corrected. For acquisition of the images in Figure 5B, a wide-angle dual speed CCD camera Sharpeye (Tröndle, Moorenweis, Germany) was used, operated by the ImageSP software. All micrographs were edited using Adobe Photoshop CS6.

**Figure 2:**
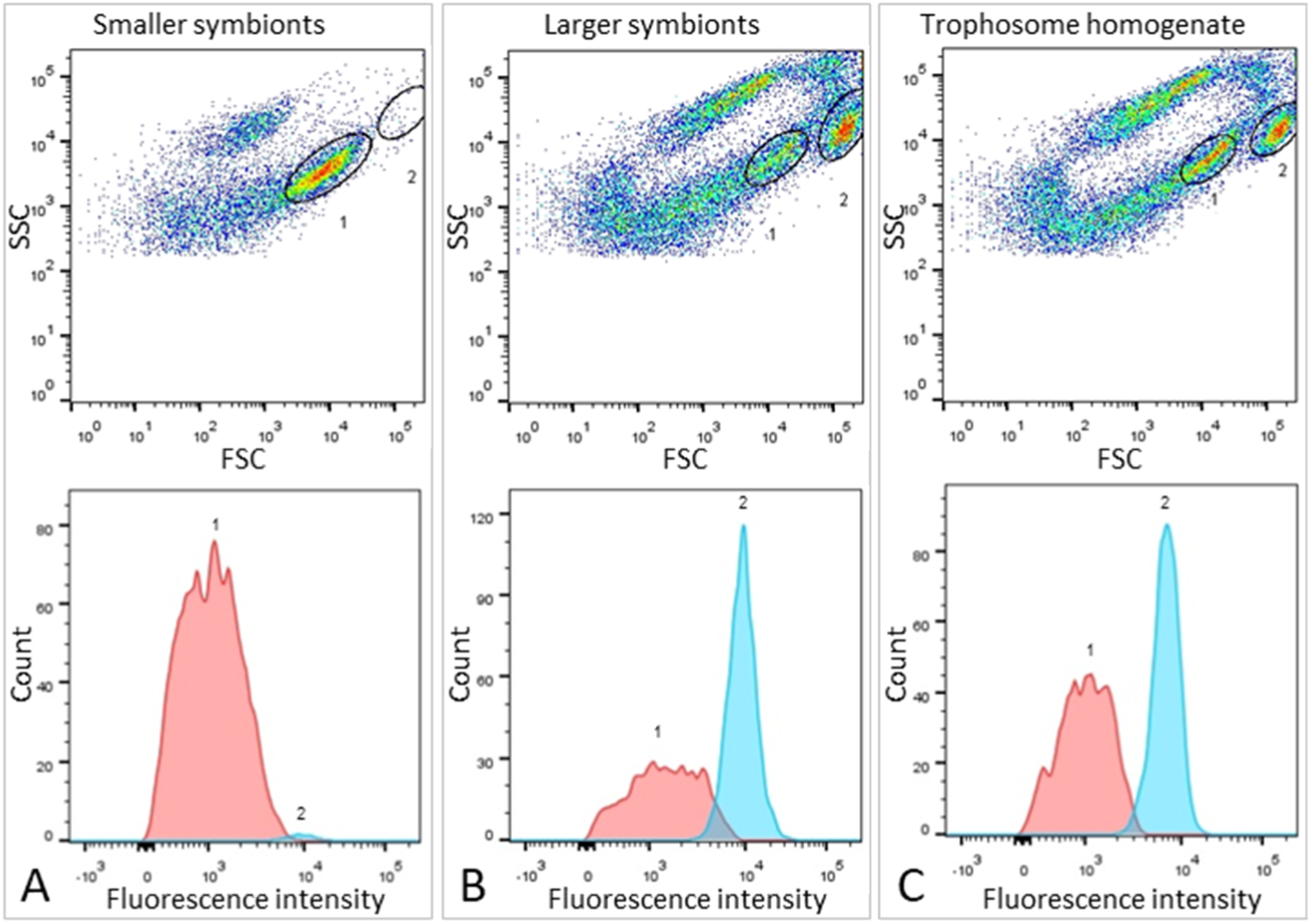
Flow cytometry for DNA quantification of *Riftia* symbionts. **A)** Dot plot of forward scatter (FSC) and side scatter (SSC), and histogram with fluorescence signal counts and fluorescence intensity per particle of a gradient fraction enriched in smaller symbionts. **B)** Gradient fraction enriched in larger symbionts. While cell population 1 was more prominent in A), population 2 was almost exclusively detected in B), and both populations were present in non-enriched trophosome homogenate (**C**), indicating that population 1 corresponds to smaller symbionts, whereas population 2 corresponds to larger symbiont cells. Cells were stained with Syto9 and median fluorescence intensity (MFI) per particle at wave length 530/30 nm was used as a measure of cellular DNA content (see Methods and Supplementary Table S8 for more details). This analysis was based on two *Rifti*a specimens with medium sulfur content.

**Figure 3:**
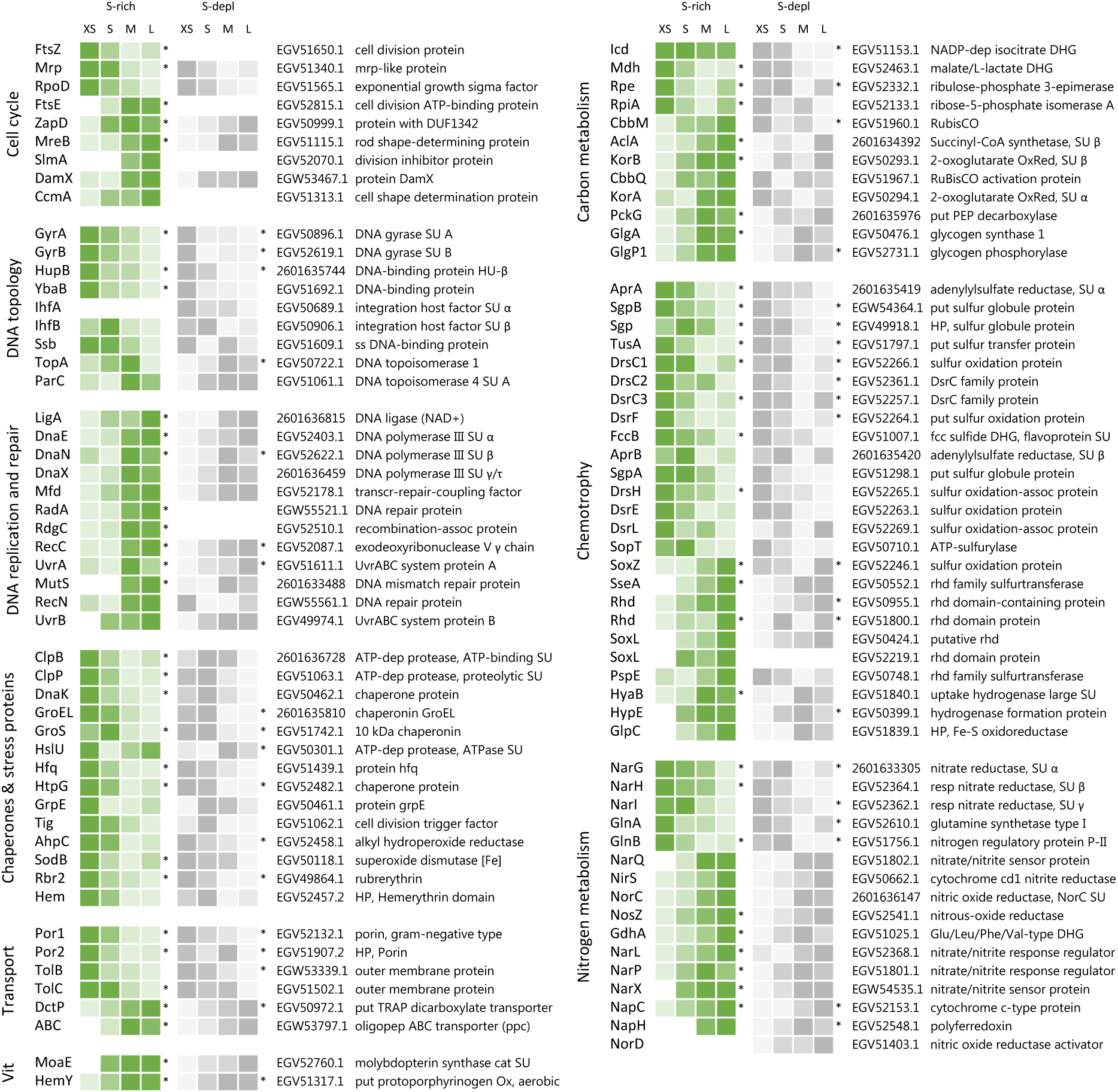
Abundance trends of selected Endoriftia proteins of various functions in the four fractions XS to L in sulfur-rich (S-rich) and sulfur-depleted (S-depl) *Riftia* specimens. Trends are indicated by color shades from light green/light grey (lowest protein abundance across all four fractions) to dark green/dark grey (highest abundance across all four fractions; note that colors do not allow comparison of protein abundance between proteins). Abundance values are based on statistical evaluation of four biological replicates (S-rich) and three biological replicates (S-depl). Proteins marked with asterisks show statistically significant trends, i.e., differences that are consistent across all replicates in S-rich or S-depl specimens (or both). White cells indicate that this protein was not detected in this sample or too low abundant to be included in statistical analyses. For an overview of all identified symbiont proteins and their relative abundances and for a summary of protein abundance trends sorted by metabolic category see Supplementary Tables S3 and S4, respectively. Accession numbers refer to NCBI/JGI entries. SU: subunit, DUF: domain of unknown function, ss: single-stranded, transcr: transcription, assoc: associated, dep: dependent, HP: hypothetical protein, put: putative, oligopep: oligopeptide, ppc: periplasmic component, DHG: dehydrogenase: RubisCO: ribulose-1.5-bisphosphate carboxylase/oxygenase, Ox: oxidase, OxRed: oxidoreductase, PEP: phosphoenolpyruvate, fcc: flavocytochrome c, rhd: rhodanese, resp: respiratory, cat: catalytic, Vit: vitamin and cofactor metabolism.

**Figure 4:**
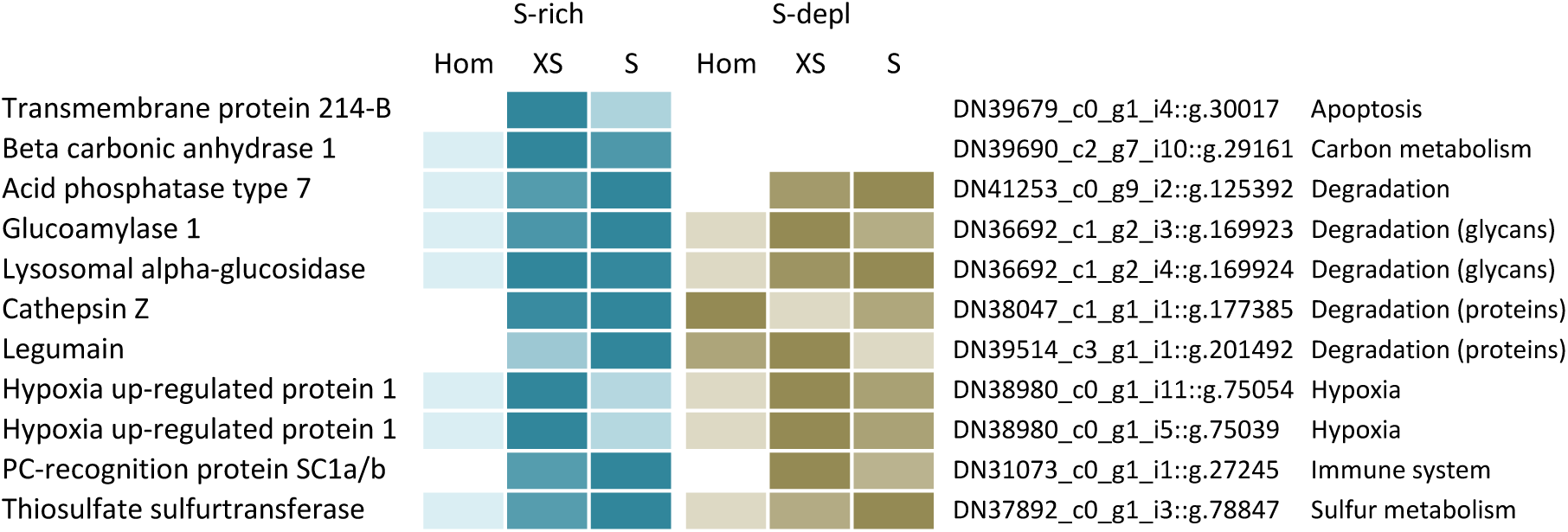
Selected *Riftia* host proteins with significantly higher relative abundance in the symbiont-enriched fractions XS and S compared to the non-enriched trophosome tissue homogenate (Hom) in sulfur-rich (S-rich) and sulfur-deleted (S-depl) *Riftia* specimens. Relative abundance trends are indicated by color shades from light blue/light brown (lowest protein abundance across the three sample types) to dark blue/dark brown (highest abundance), based on mean values from four biological replicates (S-rich) and three biological replicates (S-depl). (Note that colors do not allow comparison of protein abundance between proteins). Accession numbers refer to the combined host and symbiont database used for protein identification in this study (see Methods). For a complete list of host proteins with significantly higher abundance in fractions XS and S (compared to Hom) see Supplementary Table S5. This comparison includes only the symbiont-enriched fractions XS and S, but not fractions M and L, because these latter fractions were more likely to be contaminated by non-symbiosis-specific host proteins from host tissue fragments pelleted during centrifugation.

**Figure 5:**
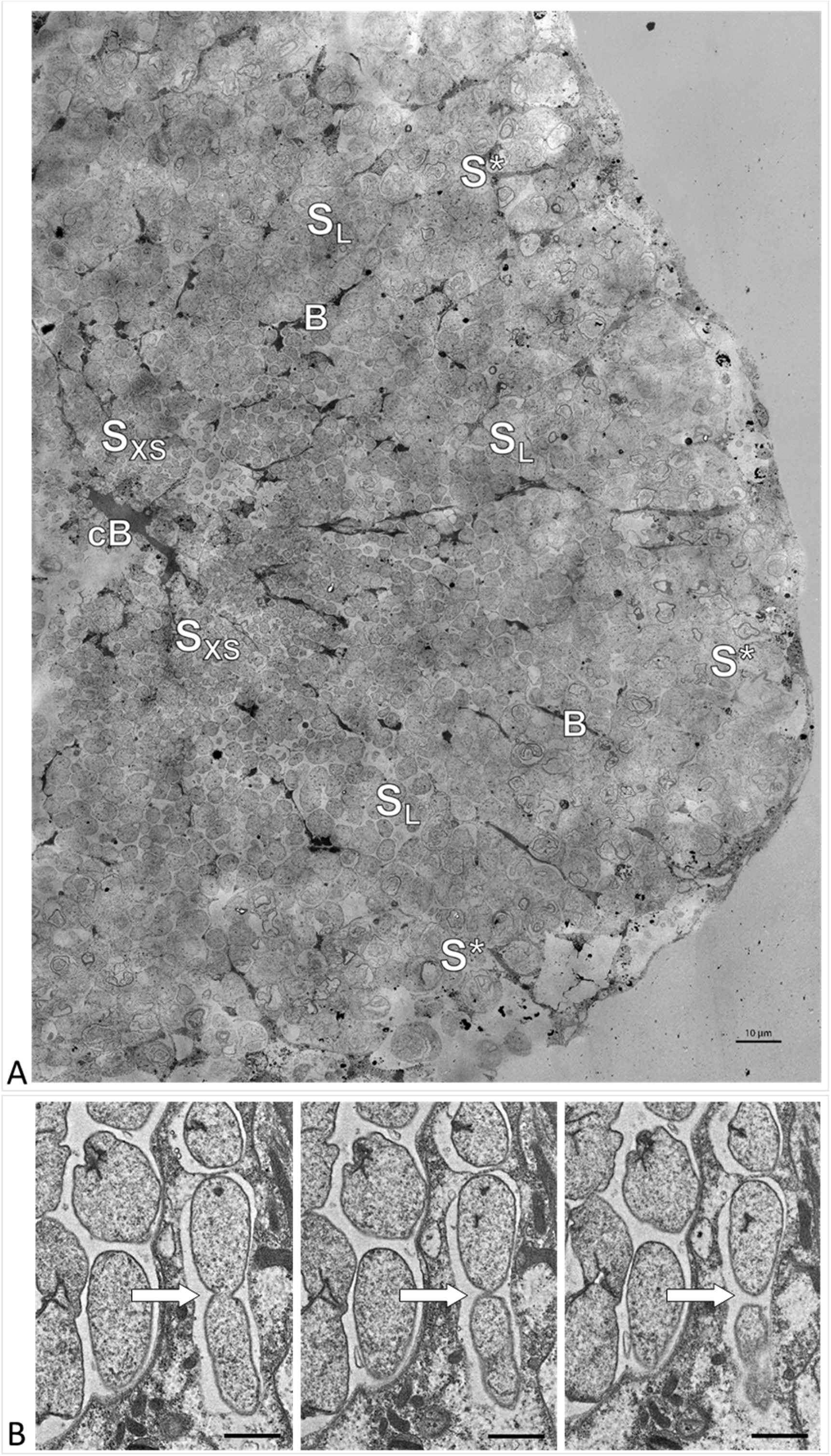
**A**) Electron micrograph of a cross section through a *Riftia* trophosome lobule. Surrounding an efferent central blood vessel (cB), small symbiont cells (S_XS_) are visible in bacteriocytes in the central lobule zone. Symbiont cell size increases towards the periphery of the lobule (S_L_: large symbiont cells). In the outermost bacteriocytes, symbiont cells are digested by host enzymes (S*). Bacteriocytes are interspersed with smaller blood vessels (B), which facilitate blood flow from the lobule periphery to the lobule center (Felbeck and Turner, 1995). The image was assembled from 50 individual transmission electron micrographs of a trophosome section from a *Riftia* specimen with sulfur-depleted trophosome. The full resolution image is available as Supplementary Figure S5. Contrast and brightness were adapted. **B**) Cell division in small *Riftia* symbionts in the trophosome lobule center of a *Riftia* specimen with sulfur-rich trophosome. All micrographs show the same dividing Endoriftia cell in three subsequent tissue sections, revealing that both daughter cells are still connected, but are about to be separated (arrow). Scale bar: 1 µm.

### HCR-FISH and confocal laser scanning microscopy (CLSM)

A gradient fraction enriched in large symbiont cells (see Supplementary Table S1 for details) that was fixed for CARD-FISH and immobilized on GTTP polycarbonate filters as described above was used for hybridization chain reaction FISH (HCR-FISH) according to Choi *et al.* (2014). We used a HCR-FISH v2.0 Custom Kit (Molecular Technologies) according to the manufacturer’s instructions. Probes targeted the *Rifti*a symbiont’s 16S rRNA (fluorescence marker: Alexa Fluor^®^ 488), and the mRNAs of ATP-citrate lyase subunit AclB (Alexa Fluor^®^ 647) and RubisCO (Alexa Fluor^®^ 594; see Supplementary Table S2 for the probe sequences). In brief, filter sections were washed twice with 50% hybridization buffer (50% formamide, 5x sodium chloride sodium citrate (SCSC, 0.75 M NaCl, 75 mM Na_3_C_6_H_5_O_7_), 9 mM citric acid, pH 6.0, 0.1% Tween 20, 50 µg/ml heparin, 1x Denhardt’s solution, 10% dextran sulfate) in 2x mPBS (89.8 mM Na_2_HPO_4_, 10.2 mM NaH_2_PO_4_, 0.9 M NaCl) at 45°C for 30 min for pre-hybridization, and incubated overnight (16 h, 45°C) with probe solution (1 pmol of each probe in 500 µl hybridization buffer). Excess probes were removed with several washing steps in 75-25% probe wash buffer (50% formamide, 5x SCSC, 9 mM citric acid, pH 6.0, 0.1% Tween 20, 50 µg/ml heparin in 5x SCSC) for 15 min at 45°C, 300 rpm, and subsequently in 5x SCSC for 30 min at 45°C and 300 rpm. Samples were pre-amplified with DNA amplification buffer (5x SCSC, 0.1% Tween 20, 10% dextran sulfate). Hairpins were activated by snap-cooling and added to the samples. After overnight incubation (16 h, room temperature) with the hairpin solution, samples were washed with 5x SCSC, containing 0.05% Tween 20 (room temperature, 300 rpm, four times 5 min, two times 30 min) and embedded in Mowiol 4-88 (Carl Roth GmbH) embedding medium prepared according to the manufacturer’s instructions. Confocal microscopy was performed on a Zeiss LSM510 meta equipped with a 100x/1.3 oil immersion objective. Probes were excited with laser lines 633 (ATP-citrate lyase), 561 (RubisCO) and 488 (16S rRNA) and signals were detected with filters suitable for dye maximal emissions at 670 nm, 595 nm and 527.5 nm, respectively. Signal intensities and cell sizes (from 8 frames showing a total of 33 cells) were quantified using the Fiji software package (Schindelin *et al.*, 2012). Individual cells were defined as regions of interest (ROI), in which signal intensity per pixel was recorded. Mean pixel intensity of ROI was calculated and background was corrected. Global background values were calculated for every channel based on up to six ROIs randomly placed in each image frame. The following cell size parameters were calculated: (i) Feret’s Diameter (the longest distance between any two points along the boundary of the ROI) and (ii) the area of the ROI.

### Flow cytometry

Subsamples of fresh homogenate, and of three gradient fractions enriched in small symbionts, and three fractions enriched in large symbionts were fixed in 1% PFA as for CARD-FISH (see above) in two biological replicates (i.e. from two *Riftia* specimens). Right before flow cytometry analysis, fixed cells were carefully pelleted and incubated in 0.1 mg/ml RNAse A (from bovine pancreas, DNase-free, Carl Roth, Germany) for 30 min at 37°C to remove RNA, and stained with Syto9 (final concentration 0.5 µmol/l in PBS), a dye that selectively stains DNA and RNA (Stocks, 2004). The fluorescence signal was analyzed using a FACSAria high-speed cell sorter (Becton Dickinson Biosciences, San Jose, CA, USA) with 488 nm excitation from a blue Coherent Sapphire solid state laser at 18 mW. Optical filters were set up to detect the emitted Syto9 fluorescence signal at 530/30nm (FITC channel). All fluorescence data were recorded at logarithmic scale with the FACSDiva 8.02 software (Becton Dickinson). Prior to measurement of experimental samples, the proper function of the instrument was determined by using the cytometer setup and tracking software module (CS&T) together with the CS&T beads (Becton Dickinson Biosciences). First, in a SSC-area *versus* FSC-area dot plot the present populations were shown. The detection thresholds and photomultiplier (PMT) voltages were adjusted by using an unstained sample. The Syto9 signal from the scatter populations was monitored in a Syto9-area histogram. For each sample at least 10.000 events in the scatter gate were recorded. For further analysis, the Syto9-stained bacteria (population 1 and 2, see Figure 2) were sorted from the bivariate dot plot, SCC-A versus Syto9 (FITC-channel). Prior to sorting, the proper function of the cell sorter was determined using the AccuDrop routine. Data analysis was done with the software FlowJo™ V10. To evaluate the results of the sorting procedure, FACS-sorted cell populations as well as unsorted subsamples of homogenate and gradient fractions were examined using an Axio Imager.M2 fluorescence microscope (Carl Zeiss Microscopy GmbH).

### Peptide sample preparation

Proteins were extracted as described in Hinzke and Markert (2017). Briefly, cells were mixed with lysis buffer (1% (w/v) sodium deoxycholate (SDC), 4% (w/v) sodium dodecyl sulfate (SDS) in 50 mM triethylammonium bicarbonate buffer (TEAB)), heated for 5 min at 95 °C and 600 rpm and cooled on ice. Samples were then placed in an ultrasonic bath for 5 min and subsequently cooled on ice. Cell debris was removed by centrifugation (14,000 x g, 10 min, room temperature). Protein concentration was determined using the Pierce BCA assay according to the manufacturer’s instructions. Peptides were generated using a 1D gel-based approach as in Ponnudurai *et al.* (2017) with minor modifications. In brief, 20 µg of protein sample was mixed with Laemmli sample buffer containing DTT (final concentration 2% (w/v) SDS, 10% glycerol, 12.5 mM DTT, 0.001% (w/v) bromophenol blue in 0.06 M Tris-HCl; Laemmli, 1970) and separated using pre-cast 4-20% polyacrylamide gels (BioRad). After staining, protein lanes were cut into 10 pieces, destained (600 rpm, 37 °C, 200 mM NH_4_HCO_3_ in 30% acetonitrile) and digested with trypsin (sequencing grade, Promega) overnight at 37 °C, before peptides were eluted in an ultrasonic bath. Peptides were then directly used for LC-MS analysis.

### LC-MS/MS analysis

MS/MS measurements were performed as described previously by Ponnudurai *et al*. (2017). In brief, samples were measured with an LTQ-Orbitrap Velos mass spectrometer (Thermo Fisher, Waltham, MA, US) coupled to an EASY-nLC II (ThermoFisher) for peptide separation using a 100 min binary gradient. MS data were acquired in data-dependent MS/MS mode for the 20 most abundant precursor ions. After a full scan in the Orbitrap analyzer (R = 30,000), ions were fragmented via CID and recorded in the LTQ analyzer.

### Protein identification and function prediction

Proteins were identified by searching the MS/MS spectra against the *Riftia* host and symbiont database (Hinzke *et al.*, 2019), which was constructed from the host transcriptome and three symbiont genome assemblies, i.e., NCBI project PRJNA60889 (endosymbiont of *Riftia pachyptila* (vent Ph05)), NCBI project PRJNA60887 (endosymbiont of *Tevnia jerichonana* (vent Tica)), and JGI IMG Gold Project Gp0016331 (endosymbiont of *Riftia pachyptila* (vent Mk28)). The cRAP database containing common laboratory contaminants (The Global Proteome Machine Organization) was added to complete the database. Database search was conducted using Proteome Discoverer v. 2.0.0.802 with the Sequest HT node as described in Kleiner *et al.* (2018) with a false discovery rate of 5% (FidoCT Protein Validator node, q-value <0.05).

To systematically screen the *Riftia* symbiont metagenome for dissimilatory sulfur metabolism-related proteins, candidates identified in different studies were searched against the *Ca.* E. persephone metaproteome database using bioedit (Hall, 1999; Supplementary Table S9). Host proteins were additionally annotated using the same tools as in Hinzke *et al.* (2019). Symbiont hydrogenase sequences were classified using HydDB (Søndergaard *et al.*, 2016).

### Statistical evaluation of metaproteomics data and abundance quantification

#### Filtering and normalization

For samples from sulfur-rich specimens, four replicates for each of the four size classes were used (resulting in 16 samples); for analysis of symbionts from sulfur-depleted specimens, three replicates were available per size class (giving a total of 12 samples). For comparisons of protein abundance (i) across different samples, e.g., to determine a protein’s abundance trend across gradient fractions XS to L, edgeR-RLE-normalized spectral counts were calculated (see below), while (ii) %orgNSAF values were used for abundance comparisons of different proteins within one sample, e.g., to determine the “most abundant” proteins in a sample.

(i) To allow for comparisons of protein abundance across different samples, spectral count data were first filtered so that they included only proteins that had at least five spectral counts in at least four out of 16 (S-rich specimens) or three out of 12 (S-depleted specimens) samples. The filtered dataset was then normalized using Relative Log Expression (RLE) normalization with the package edgeR v.3.24.3 (Robinson *et al.*, 2010) in R v. 3.5.1 (R Core Team, 2018; Supplementary Table S3a). The filtering and normalization step was included to avoid biasing the analysis towards symbiont proteins that were only detected in the high-density fractions M and L (enriched in larger symbiont cells), but which were absent in fractions of lower density (XS and S, containing primarily smaller cells). Fractions S and particularly XS contained relatively more host proteins, leading to a lower total number of detectable symbiont proteins. (Note that these values were not normalized to protein size, so that a protein’s relative abundance changes can be followed across different samples, but abundances cannot be compared between proteins). We tested for significant differences in symbiont protein abundance between individual gradient factions (representing enrichments of different cell size classes) using two methods, i.e., hierarchical and profile clustering and random forests (see below).

(ii) To be able to compare relative symbiont protein abundances within samples and to identify particularly abundant proteins, %orgNSAF values were calculated from unfiltered spectral counts by normalization to protein size and to the sum of all proteins in a sample (Zybailov *et al.*, 2006, Mueller *et al.*, 2010). %orgNSAF values give an individual protein’s percentage of all proteins in the same sample (Supplementary Table S3b). Note that %orgNSAF values cannot be compared across different samples, due to the unequal number of total host and symbiont proteins in different samples.

#### STEM analysis

For protein expression profile clustering, we employed the Short Time Series Expression Miner (STEM; Ernst and Bar-Joseph, 2006) v. 1.3.11., which fits gene expression profiles in ordered short series datasets (like the cell cycle stages of *Ca.* E. persephone), to model profiles representing different expression patterns. Filtered and RLE-normalized data were log-normalized, repeat data were defined to be from different time points and data were clustered using the STEM method with default options. For STEM filtering, the minimum correlation between repeats and the minimum absolute expression change were set to 0.5. All permutations were used. For correction, the false discovery rate (FDR) was set to 0.05. Profiles were clustered with a minimum correlation percentile of 0.5. Other parameters were left at default values. Proteins which were assigned to model profiles, i.e., all proteins which were not removed by filtering and showed a consistent trend in all replicates, were used for further analysis. This means that differences in protein abundance patterns were considered significant if proteins were detected with a consistent abundance trend across all replicates (increase, decrease or alternating increase and decrease of abundance from fraction XS to L).

#### Random forests

For random forest analysis, we used the ranger package v. 0.10.1 (Wright and Ziegler, 2015) in R v. 3.5.1 (R Core Team, 2018). Random forests are a machine learning technique, which can be used to find the variables – here proteins – that allow to predict which datasets or samples are similar (and which ones are not; Degenhardt *et al.*, 2019). For variable importance calculation, we employed the method from (Janitza *et al.*, 2018) as implemented in the ranger package. This method uses a heuristic approach, where a null distribution for p-value calculation is generated based on variables with importance scores of zero or negative importance scores. For pairwise comparisons, the data set was subjected to an additional filtering step, so that only proteins with a minimum of five spectral counts in at least six out of eight (S-rich) or four out of six (S-depleted) samples were included. The comparison of all 16 S-rich samples included only such proteins which had a minimum of five spectral counts in at least five samples, and the comparison of all 12 S-depleted samples included only proteins with five or more spectral counts in a minimum of four samples. The filtered and RLE-normalized data were used for random forest analysis as follows: 2,000 forests with 10,000 trees per forest were grown for pairwise comparisons as well as for comparisons including the samples representing all four size classes. Proteins which had a p-value below 0.05 in >90% of the forests were included in further analyses.

#### Significant differences in protein abundance

Proteins that showed significant abundance differences as determined by STEM analysis or by random forest analysis (see above) or by both methods were included in a common list. Please note that this approach of determining significant protein abundance differences was not based on individual p-values. For proteins with significant abundance differences, we clustered the z-scored mean abundances using hierarchical clustering (Pearson correlation, complete linkage) in R to visualize their abundance trends (Supplementary Figure S1). For this purpose, we employed the R base package stats (R Core Team, 2018) as well as the packages cluster (Maechler *et al.*, 2018) and ComplexHeatmap (Gu *et al.*, 2016). For comparison of S-rich and S-depleted symbionts of the same size class, we used the R package edgeR v. 3.24.3 (Robinson *et al.*, 2010), which uses a Bayes-moderated Poisson model for count data analysis, with an overdispersion-adapted analogon to Fisher’s exact test for detecting differentially expressed genes (Robinson *et al.*, 2010).

#### Host proteins

Host proteins which were more abundant in symbiont-enriched fractions as compared to the non-enriched trophosome homogenate are candidates for direct host-symbiont interaction, as they might be secreted into symbiont compartments or even physically associated with symbiont cells. For evaluation of host protein enrichment, we used fractions XS and S enriched in the two smallest symbiont size classes (i.e. fractions collected from the upper part of the gradient). As the larger gradient fractions sometimes contained the gradient pellet, in which host proteins can also accumulate when host tissue fragments are pelleted, these fractions were not used for host protein analysis. Comparisons of relative host protein abundance between trophosome homogenate and fractions XS and S were performed using the R package edgeR v. 3.24.3. Spectral count data were filtered to include only proteins which had at least five spectral counts in at least four (for S-rich specimens) or three (in S-depleted specimens) samples and RLE-normalized abundance values were compared between samples. Proteins which were significant in the edgeR comparison and had a higher mean RLE-normalized abundance in fractions XS and S than in the homogenate sample were included in functional analysis.

### Data availability

The mass spectrometry proteomics data have been deposited to the ProteomeXchange Consortium (ProteomeXchange – ProteomeCentral) via the PRIDE partner repository (Vizcaíno *et al.*, 2016) with the dataset identifier PXD016986. The dataset will be released upon acceptance of the manuscript in a peer-reviewed journal.

## Results

### Enrichment of individual symbiont cell sizes by gradient centrifugation

Our rate-zonal gradient centrifugation approach allowed us to enrich distinct symbiont cell sizes from *Riftia* trophosome tissue. Based on CARD-FISH microscopy, we defined four size ranges (Figure 1): very small symbiont cells (≥2.0 – <3.9 µm diameter), small (≥3.9 – <5.3 µm), medium (≥5.3 – <6.8 µm) and large symbiont cells (≥6.8 – 20.0 µm; see also Supplementary Results and Discussion A). For subsequent comparative metaproteomic analyses, we chose the gradient fractions that were most enriched in one of these cell size ranges. In the following, these four gradient fractions are referred to as XS (containing the highest percentage of very small symbiont cells) to L (containing the highest percentage of large cells). The enrichment procedure was highly reproducible, particularly for symbionts isolated from sulfur-rich trophosome tissue (Figure 1).

### Symbiont DNA quantification

Flow cytometry and fluorescence-activated cell sorting (FACS) indicated that DNA content in large *Riftia* symbionts is up to 10-fold higher compared to small symbionts. To identify distinguishable bacterial cell populations, we examined Syto9-stained cells in *Riftia* trophosome homogenate and in gradient fractions from the upper and lower parts of the gradient (enriched in smaller and larger symbionts, respectively) with regard to their light scattering properties. Forward scatter (FSC) and side scatter (SSC) usually correlate with cell size and cell granularity, respectively (Bouvier *et al.*, 2001, Tracy *et al.*, 2010). Amongst a number of particle groups with different properties (see also Supplementary Results and Discussion C), we found two populations, 1 and 2, which were abundantly detected in non-enriched trophosome homogenate, but showed very dissimilar frequencies in fractions enriched in larger or smaller symbionts (Figure 2, Supplementary Figure S2): While population 1, which exhibited relatively lower FSC and SSC signals (indicative of smaller cell size and lower cell complexity), was highly abundant in fractions enriched in smaller symbionts, this population was notably less prominent in fractions enriched in larger symbionts. Simultaneously, population 2, which gave higher FSC and SSC signals (indicative of larger cell size and higher complexity), was highly abundant in fractions enriched in larger symbionts but nearly absent in gradient fractions enriched in smaller symbionts. This suggests that populations 1 and 2 consist of smaller and larger symbionts, respectively. This assumption was verified by FACS-separation of both populations from trophosome homogenate, and examination of the sorted cell suspensions by fluorescence microscopy along with unsorted enriched gradient fractions and homogenate samples for reference (Supplementary Figure S2). For quantification of DNA in smaller and larger symbionts, we compared median fluorescence intensities (MFI) per particle between population 1 and 2 in non-enriched homogenate and in enriched gradient fractions. In all sample types, MFI per particle was notably lower in population 1 (between 186 and 1,994 relative fluorescence units, rfu) than in population 2 (2,712 – 10,723 rfu). On average, MFI was 9.7-fold higher in population 2 than in population 1 (Supplementary Table S8).

### Protein identifications and relative protein abundance

We identified a total of 1,946 symbiont proteins across all sample types, including the four gradient fractions XS – L and non-enriched homogenate from both, sulfur-rich and sulfur-depleted *Riftia* specimens (Supplementary Table S6). Our sample fractionation by gradient centrifugation thus facilitated detection of around 60% of the symbiont’s theoretical proteome, which encompasses 3,182 proteins in PRJNA60889, and yielded substantially higher symbiont protein identification rates than non-enriched trophosome homogenate samples alone (1,223 total symbiont protein identifications). After stringent filtering and normalization, a subset of 1,212 symbiont proteins from gradient fractions XS – L was included in statistical analysis using abundance profile clustering and random forests (Supplementary Table S3). A total of 465 proteins showed significant differences in relative abundance in S-rich and/or S-depleted samples (Supplementary Figure S1; note that the term “significant” denominates trends that were consistent across all replicates in the context of our statistical approach). In Figure 3 and Supplementary Table S3, proteins that showed such significant changes in relative abundance are marked with asterisks.

Of all proteins with significant abundance changes, 56% (261 proteins) followed a clear, continuous abundance trend from fraction XS to L or *vice versa*, that is, protein abundance increased or decreased with increasing symbiont cell size (Supplementary Table S3). For the majority of symbiont proteins, abundance trends in samples obtained from sulfur-rich (energy-rich) and sulfur-depleted (energy-depleted) trophosome tissue were highly similar. Very few proteins were detected only in sulfur-rich samples (61 of 1,212 proteins) or exclusively in sulfur-depleted samples (77 proteins). For a discussion of specific differences observed between symbionts from energy-rich and energy-starved trophosome tissue, see Supplementary Results and Discussion B.

### Symbiont protein functions

#### Cell cycle, DNA topology, replication and repair

Proteins involved in the bacterial cell cycle and in DNA topology, -replication and -repair were differentially expressed across fractions XS to L (Figure 3, Supplementary Table S4a). While the cell division protein FtsZ, DNA gyrase and DNA-binding proteins decreased significantly in abundance from fraction XS to L, abundance of other cell division-related proteins (e.g., FtsE, MreB, division inhibitor SlmA), and of proteins involved in DNA replication (e.g., DNA ligase, DNA polymerase) and repair (e.g., UvrAB) increased. Interestingly, FtsZ abundance was very low in S-depleted fractions, so that it was excluded from statistical analysis in these samples (see Supplementary Results and Discussion).

#### Chaperones and stress proteins

Many chaperones and other proteins involved in protein folding, as well as oxidative stress-related proteins were detected with significantly decreasing abundance from fraction XS to L, including (amongst others) the proteases ClpB, ClpP, GroEL, the abundant alkyl hydroperoxide reductase AhpC, superoxide dismutase SodB, and rubrerythrin (Figure 3, Supplementary Table S4b).

#### Transport

Outer membrane proteins such as two porins and TolBC showed significant abundance differences between the fractions, with highest relative abundance in fraction XS and lowest abundance in fractions L. Porin EGV52132.1 (Por1) was the most abundant symbiont protein throughout all sample types (Figure 3, Supplementary Table S4c). On the other hand, all five detected tripartite ATP-independent periplasmic (TRAP) transporter subunits and ten out of 13 ABC transporter components were relatively more abundant in fraction L (see also Supplementary Results and Discussion D).

#### Central metabolism

##### Carbon metabolism

Several tricarboxylic acid (TCA) cycle enzymes (e.g., Icd, Mdh), as well as enzymes of the pentose phosphate pathway (e.g., Rpe, RpiA) were detected with decreasing abundances from fraction XS to L (Figure 3, Supplementary Table S4d). In contrast, the key enzymes of the two CO_2_-fixing pathways, Calvin cycle (RubisCO, CbbM) and rTCA cycle (ATP-citrate lyase, AclA; oxoglutarate oxidoreductase, KorAB), as well as most of the gluconeogenesis-related (e.g., PckG), and glycogen metabolism-related enzymes (e.g., GlgA, GlgP) increased in abundance from fraction XS to L.

##### Chemotrophy

Many sulfide oxidation-specific proteins, including both subunits of the abundant key enzyme adenylylsulfate reductase AprAB, as well as proteins involved in sulfur storage (sulfur globule proteins) had their highest abundance in fraction XS or S and their lowest abundance in fraction M or L (Figure 3, Supplementary Table S4e, Supplementary Figure S4). In contrast, thiosulfate oxidation-related proteins like SoxZ, SoxL and other rhodanese-like proteins were detected with significantly increasing abundance from fraction XS to fraction L. Four additional Sox proteins, i.e., SoxA, SoxB, SoxW and SoxY, which were detected at very low abundances across the sample types (and were therefore excluded from statistical analysis), were identified in fraction M and L, but were completely absent from fraction XS (Supplementary Table S3). Three proteins involved in energy generation by hydrogen oxidation, HyaB, HypE and GlpC, were also detected with increasing abundance from fraction XS to fraction L.

##### Nitrogen metabolism

Relative abundance of all three respiratory membrane-bound nitrate reductase subunits, NarGHI, decreased significantly from fraction XS to L, as did abundance of glutamine synthetase GlnA (Figure 3, Supplementary Table S4f, Supplementary Results and Discussion E). On the other hand, various other denitrification-related proteins (such as nitrite reductase NirS, nitrous oxide reductase NosZ, and nitrate/nitrite signal transduction systems) and glutamate dehydrogenase GdhA showed relatively higher abundances in fraction L (or M) than in fraction XS. The same trend was observed for the periplasmic nitrate reductase components NapC and NapH. Moreover, NapG, another NapH copy, the nitric oxide reductase subunit NorB, nitric oxide reductase activation protein NorQ, and the putative assimilatory nitrite reductase subunit NirB, whose overall abundances were too low to include them in statistical analysis, were only detected in fraction M and/or L.

#### Other categories

50 (75%) of the 67 proteins involved in cofactor- and vitamin synthesis in S-rich samples had their highest abundance in fraction M or L (Figure 3, Supplementary Table S4g). Also, of the 33 identified tRNA ligases and tRNA synthetases, 25 (75%) were most abundant in fraction M or L (in S-rich samples, Supplementary Table S4h).

### Symbiosis-specific host proteins

Our density gradient fractionation procedure allowed not only for the identification of symbiont proteins with differential abundance across different Endoriftia size ranges, but also enabled us to single out host proteins that are potentially involved in direct interactions with the symbionts. As host proteins that are attached to the symbionts are pulled down with the symbiont cells during gradient centrifugation, these proteins should be significantly more abundant in symbiont-enriched fractions compared to the non-enriched trophosome homogenate (Figure 4, Supplementary Table S5). Besides many ribosomal and mitochondrial host proteins, which were also enriched, putatively symbiont-associated host proteins included the host’s peptidoglycan-recognition protein SC1a/b, beta carbonic anhydrase 1, digestive proteins involved in protein- and carbohydrate degradation, e.g. acid phosphatase, digestive proteases and glycan degradation enzymes, as well as hypoxia up-regulated proteins, a thiosulfate sulfurtransferase, and transmembrane protein 214-B.

## Discussion

### Symbiont growth and differentiation

#### Cell division plays a more prominent role in small symbionts

As indicated by the significant decrease in abundance of the cell division key protein FtsZ from fraction XS to fraction L, small Endoriftia are more engaged in cell division than larger symbionts. In accordance with the microscopy-based hypothesis of Bright and Sorgo (2003), the smallest symbionts, which are *in situ* localized in the trophosome lobule center (Figure 5), thus apparently function as stem cells of the symbiont population. During cell division, FtsZ forms the Z ring, to which the other division-related proteins are successively recruited (reviewed in Weiss, 2004). Cell size and cell division therefore likely depend on the amount of FtsZ available (Chien *et al.*, 2012). This correlation is, for example, also reflected by a decrease of FtsZ concentration during differentiation of vegetative cells into non-dividing larger heterocysts in the cyanobacterium *Anabaena* (Klint *et al.*, 2007). Interestingly, while FtsZ abundance decreased across fractions, many other proteins which interact with FtsZ during cell division were detected with increasing abundance from fraction XS to L. This indicates that these proteins are also involved in processes other than cell division, e.g., in determining cell shape and stabilization. ZapD, for example, is involved in FtsZ filament organization, and its overexpression leads to cell filamentation (Durand-Heredia *et al.*, 2012). DamX overexpression, too, was observed to induce filamentation in *E. coli* (Lyngstadaas *et al.*, 1995), while overexpression of the cell shape determination protein CcmA in *E. coli* and *P. mirabilis* lead to enlarged, ellipsoidal cells (Hay *et al.*, 1999), and FtsEX is required for cell elongation rather than cell division in *B. subtilis* (Domínguez-Cuevas *et al.*, 2013). The actin homolog MreB is pivotal for rod-shape formation in bacteria and for cell stiffness in *E. coli*, could negatively regulate cell division, and participates in chromosome segregation (Wachi and Matsuhashi, 1989, Kruse *et al.*, 2006, Wang *et al.*, 2010, reviewed in Reimold *et al.*, 2013). In large Endoriftia, these proteins might therefore be involved in stabilizing growing symbiont cells. SlmA, which was only detected in fractions M and L in our study, was shown to disassemble FtsZ polymers, thus acting as a cell division inhibitor (Cho *et al.*, 2011), which supports the idea of relatively less cell division in large *Riftia* symbionts. Although Endoriftia’s major cell division protein FtsZ was notably (1.75x) less abundant in fraction L (compared to fraction XS), it was not completely absent. This may indicate that cell division is reduced with increasing cell size, but not abandoned altogether, or it may point to additional FtsZ functions, besides cell division (as also suggested for *Anabeana* (Klint *et al.*, 2007) and *E. coli* (Thanedar and Margolin, 2004)).

#### Large symbionts have more genome copies and less compact chromosomes

Endoriftia’s differentiation into large, non-dividing (but still replicating) cells coincides with endoreduplication cycles and an increase in genome copy number, as indicated by our flow cytometry analysis (Figure 2, Supplementary Figure S2, Supplementary Table S8). This observation is in agreement with earlier findings of Bright and Sorgo (2003), who noted more than one chromatin strand-containing area in large coccoid *Riftia* symbiont cells in electron microscopy images, whereas small rods and cocci featured only one chromatin strand area. The idea of endoreduplication in larger *Riftia* symbionts is also supported by the observation that large symbiont cells, which apparently divide less frequently than smaller cells (see above), still actively replicate DNA, as indicated by high abundances of DNA ligase and DNA polymerase III in fraction L. The observed decreasing abundance of DNA gyrase GyrAB with increasing cell size additionally corroborates this idea, as type II topoisomerases such as gyrase are not only involved in supercoiling and initiation of DNA replication (Levine *et al.*, 1998, Nöllmann *et al.*, 2007), but also essential for decatenation of newly replicated chromosomes in bacteria (Steck and Drlica, 1984, Guha *et al.*, 2018). Moreover, inhibition of topoisomerase II in eukaryotes leads to endoreduplication and polyploidy (Cortés and Pastor, 2003, Cortés *et al.*, 2003). Polyploidy in thiotrophic symbionts was also observed in the lucinid bivalve *Codakia orbicularis*, where larger symbiont cells contained more than four genome copies, while smaller cells had only one genome copy (Caro *et al.*, 2007), and in ectosymbionts of *Eubostrichus* nematodes, in which up to 16 nucleoids per large symbiont cell were reported (Polz *et al.*, 1992, Pende *et al.*, 2014). Moreover, also terminally differentiating *Rhizobia* undergo endoreduplication cycles (Mergaert *et al.*, 2006), and high genome copy numbers have been reported for various bacterial insect symbionts, e.g., of aphids, cockroaches and sharpshooters (Komaki and Ishikawa, 2000, López-Sánchez *et al.*, 2008, Woyke *et al.*, 2010), suggesting that polyploidy is common in symbiotic bacteria. Possibly, enlarged polyploid cells might increase the metabolic activity and/or fitness of the Endoriftia cells: In *E. coli*, a *mreB* point mutation led to increased cell size, which gave the cells a measurable fitness advantage in presence of certain carbon sources (Monds *et al.*, 2014). Moreover, polyploidy was suggested to provide evolutionary advantages like a low mutation rate and resistance towards DNA-damaging conditions in haloarchaea (Zerulla and Soppa, 2014). In plants, endoreduplication is common and might increase transcription and metabolic activity of the cells (Kondorosi and Kondorosi, 2004), leading to enhanced productivity (Sattler *et al.*, 2016). More generally, in symbiotic associations, where the bacteria are stably and sufficiently provided with carbon and energy sources, the advantages of polyploidy might be greater than the associated costs (Angert, 2012).

Higher genome copy numbers in large symbionts seem to be accompanied by a lower degree of DNA condensation, compared to small Endoriftia, as indicated by notably lower abundances of the histone-like DNA-binding proteins HU (HupB) and integration host factor (IHF, IhfAB), and of DNA gyrase GyrAB in fraction L, compared to XS. Bacterial histone-like DNA-binding proteins like HU and IHF structure the chromosome and modulate the degree of supercoiling (reviewed in Dorman and Deighan, 2003). In *E. coli*, absence of HU leads to unfolding of the chromosome and cell filamentation (Dri *et al.*, 1991), and unspecific DNA-binding by IHF was shown to contribute to DNA compaction (Ali *et al.*, 2001). Moreover, bacterial DNA gyrase was also suggested to be involved in nucleoid compaction in *E. coli* (Stuger *et al.*, 2002). Co-occurrence of endoreduplication and decondensated DNA is also known in plant cells (Kondorosi and Kondorosi, 2004). As decondensation occurs in actively transcribed DNA regions (Wang *et al.*, 2014), it might facilitate protein synthesis and metabolic activity in large Endoriftia.

Since DNA condensation may function as a DNA protection mechanism (Ohniwa *et al.*, 2006, Mukherjee *et al.*, 2008, Yoshikawa *et al.*, 2008, Takata *et al.*, 2013), less condensed DNA might be more prone to various kinds of damage and require the enhanced expression of DNA repair mechanisms. This would explain the observed higher abundance of several DNA repair proteins in fraction L, which was enriched in larger, older symbiont cells with (presumably) larger quantities of less condensed DNA, compared to the smaller symbiont cells. RadA, RdgC, RecCN, UvrAB, and Mfd, which are known to be involved in DNA recombination and repair in many bacteria (Kowalczykowski, 2000, Beam *et al.*, 2002, Tessmer *et al.*, 2005, Drees *et al.*, 2006, Truglio *et al.*, 2006, Deaconescu *et al.*, 2007), may compensate for this elevated vulnerability. In eukaryotes, chromatin decondensation was shown to facilitate access of the DNA damage response to double strand breaks, thus allowing for more efficient repair (Murga *et al.*, 2007).

#### Small symbionts may be exposed to elevated stress levels

Small symbionts might experience cell division-related or host-induced stress in the early phase of their cell cycle, as indicated by elevated levels of symbiont chaperones and stress response proteins, as well as of reactive oxygen species (ROS) scavengers in fraction XS. This is in line with observations in *Caulobacter crescendus*, where the DnaK-DnaJ and GroEL-GroES systems are crucial for cell division (Susin *et al.*, 2006), and in *E. coli*, where the protease ClpXP and the RNA chaperone Hfq are probably involved in cell division as well (Camberg *et al.*, 2009, Zambrano *et al.*, 2009). Interestingly, like the putative Endoriftia stem cells, eukaryotic embryonic stem cells also feature high levels of chaperone expression and stress tolerance (Prinsloo *et al.*, 2009). Although the reason for this congruence is yet unknown, possibly, cell division-related processes might require elevated levels of chaperones and stress proteins, e.g. to ensure correct assembly of all parts of the division machinery or to counteract some sort of yet to be determined host-induced stress.

Possibly, such host-induced stress may also involve the production of ROS in symbiont-containing bacteriocytes, similar to animal and plant hosts, which generate ROS to defend themselves against pathogenic bacteria (Heath, 2000, Lynch and Kuramitsu, 2000, D’Haeze and Holsters, 2004). Small symbionts, which are relatively loosely packed in their host cell vesicles (Figure 5A) and have a comparatively high surface-to-volume ratio, might be particularly exposed to this presumptive ROS stress, while larger symbionts, which are more tightly packed, may face lower ROS levels. This would explain the observed higher abundance of the ROS scavengers rubrerythrin (Rbr2), superoxide dismutase (SodB), and alkylhydroperoxide reductase (AhpC) in small symbionts. In line with this assumption, a superoxide dismutase and also the chaperones ClpB, HtpG, and DnaK were suggested to be involved in ROS protection in *Serratia symbiotica* (Renoz *et al.*, 2017), and ClpB protease expression has been shown to increase during oxidative stress in the intracellular pathogen *Francisella tularensis* (Twine *et al.*, 2006).

Interestingly, neither S-depleted nor S-rich samples showed indications of a strong bacterial stress response in fraction L, indicating that imminent digestion by the host poses no particular stress to the large symbionts. Possibly, bacterial degradation happens too fast to elicit a stress response, or a stress response is suppressed during symbiosis, either by the symbionts themselves or by the host via a yet to be determined mechanism.

#### Host-microbe interactions may be particularly important in small Endoriftia

Abundant Endoriftia membrane proteins might play a key role in host interaction in small symbionts. Particularly, the high and differential abundance of porin Sym EGV52132.1, the most abundant symbiont protein in all fractions, which was nearly 3-times more abundant in fraction XS (11.7 %orgNSAF) than in fraction L (4.0 %orgNSAF), suggests that this protein may be of varying relative importance throughout the symbiont’s differentiation process. Porins are water-filled channels in the outer membrane, through which small hydrophilic molecules can diffuse (Fernández and Hancock, 2012). In the oyster pathogen *Vibrio splendidus*, the porin OmpU serves as adhesin or invasin and is involved in recognition by the host cell (Duperthuy *et al.*, 2011), while in *Neisseria gonorrhoeae*, a porin inhibits phagocytosis by human immune cells (Mosleh *et al.*, 1998, Lorenzen *et al.*, 2000). Interestingly, the phagocytosis-inhibiting action of *N. gonorrhoeae* porin apparently involves interference with the host’s oxidative burst, i.e., the porin allows the pathogen to evade killing by host-produced ROS (Lorenzen *et al.*, 2000). Although the exact function of Endoriftia porin has not been elucidated yet, we suggest that it may have a similar function in resistance against host stress or ROS. This would be in line with elevated levels of ROS scavengers in small *Riftia* symbionts (see above). Porins are furthermore not only known to be involved in recognition by the host (e.g., in the squid symbiont *Vibrio fischeri* (Nyholm *et al.*, 2009)), but were also shown to be involved in survival in and communication with the host in other intracellular and pathogenic bacteria, rendering *Vibrio cholerae* and *Xenorhabdus nematophila* more resistant against antimicrobial compounds (Mathur and Waldor, 2004, van der Hoeven and Forst, 2009). As *Riftia* trophosome tissue has antimicrobial effects (Klose *et al.*, 2016), and considering that *Riftia* might employ histone-derived antimicrobial peptides to modulate the symbiont’s cell division (Hinzke *et al.*, 2019), Endoriftia porin may enable the symbionts to reject antimicrobial compounds produced by the host. This would be of particular importance for small symbionts, as it would ensure survival of the symbiont stem cell subpopulation and sustain their division capability.

Besides porin, the symbiont’s outer membrane efflux pump TolC was also most abundant in fraction XS, suggesting that it may play a similar role in host interaction or persistence. TolC is a versatile export protein of Gram-negative bacteria, which interacts with different transporters of the cytoplasmic membrane to export proteins and drugs (reviewed in Koronakis *et al.*, 2004). In *Sinorhizobium meliloti*, TolC is apparently involved in establishing the symbiosis with legumes, possibly by conferring increased stress resistance and by secreting symbiosis factors (Cosme *et al.*, 2008), while *Erwinia chrysanthemi* TolC enables re-emission of the antimicrobial compound berberine and is thus essential for *Erwinia* growth in plant hosts (Barabote *et al.*, 2003).

Microbe-host interactions with particular relevance in smaller Endoriftia may furthermore also be mediated by chaperones and stress proteins, which were most abundant in fraction XS (see above). Chaperones have been shown to play a role in host interaction and intracellular survival in several pathogenic and symbiotic bacteria. For example, DnaK appears to be essential for growth of *Brucella suis* in phagocytes (Köhler *et al.*, 1996), while HtpG seems to be involved in virulence and intracellular survival of *Leptospira* (King *et al.*, 2014), *Salmonella* (Verbrugghe *et al.*, 2015) and *Edwardsiella tarda* (Dang *et al.*, 2011). Mutations in the post-transcriptional regulator *hfq* often lead to reduced fitness and virulence in bacterial pathogens (reviewed in Chao and Vogel, 2010). Moreover, ClpB in *Listeria* is apparently specifically involved in virulence (Chastanet *et al.*, 2004), as are ClpX and ClpP in *Staphylococcus aureus* (Frees *et al.*, 2003). In the insect symbiont *Wolbachia*, HU beta was suggested to directly interact with the host (Beckmann *et al.*, 2013). Additional symbiont proteins that may protect small *Endoriftia* from host interference, and particularly so in S-depleted *Riftia* specimens, included an ankyrin protein and an FK506-binding protein (see Supplementary Results and Discussion B).

#### Interaction-specific host proteins

We detected a number of ‘symbiosis-specific’ *Riftia* proteins, which were co-enriched with symbiont cells in fractions XS and/or S and may thus facilitate direct host-microbe interactions or enable the host to provide optimal conditions for the symbiont. PGRPs, for example, are involved in innate immunity (Kang *et al.*, 1998) and have previously been shown or suggested to participate in symbiotic interactions (Troll *et al.*, 2009, Wang *et al.*, 2009, Royet *et al.*, 2011, Wippler *et al.*, 2016). Since oxygen concentrations in the trophosome might be comparatively low (benefitting the microaerophilic symbionts; Hinzke *et al.*, 2019), the hypoxia up-regulated *Riftia* proteins we detected may present a protective adaptation of the host to these hypoxic conditions. In support of this idea, Hyou1 was shown to have a protective function during hypoxia in human cells (Ozawa *et al.*, 1999). Moreover, enrichment of beta carbonic anhydrase 1, which interconverts bicarbonate and CO_2_, suggests that this host protein serves to optimally provide the symbionts with CO_2_ for fixation. The host transmembrane protein 214-B (TMP214-B), which was exclusively detected in symbiont-enriched fractions (but not in trophosome homogenate) may be involved in cell death of symbiont-containing bacteriocytes by an apoptosis-related mechanism. This would be in line with our previous suggestion that apoptosis-related proteins may play a role in symbiont and bacteriocyte cell death (Hinzke *et al.*, 2019), and is further supported by the fact that TMP214-B was shown to be involved in apoptosis caused by endoplasmic reticulum stress (Li *et al.*, 2013). The detection of degradation proteins such as cathepsin Z, legumain, glucoamylase 1 and lysosomal alpha-glucosidase in fractions XS and S furthermore implies that the host digests not only large symbiont cells in the degradative trophosome lobule zone (see Figure 5A), but that small symbionts might also be exposed to host digestion.

### Metabolic diversity among symbiont size classes

#### Large symbionts focus on carbon fixation and biosynthesis

Highest individual abundances of various carbon fixation and biosynthesis-related enzymes as well as highest overall abundances of all biosynthetic categories (including carbon-, amino acid-, lipid-, nitrogen- and cofactor metabolism; Supplementary Table S7) in fraction L suggests that large Endoriftia cells are relatively more engaged in the production of organic material than smaller symbiont cells. In support of this idea, we observed notably higher RubisCO mRNA signal intensity in large symbiont cells than in smaller *Riftia* symbionts in our HCR-FISH analysis (Supplementary Results and Discussion D, Supplementary Figure S3). This concurs with an autoradiographic study of Bright *et al*. (2000), who observed highest ^14^C carbon incorporation in the *Riftia* trophosome lobule periphery and lowest short-term incorporation in the lobule center. As previously suggested (Hand, 1987), these and other observed differences might be due to a biochemical gradient, which could be caused by the direction of blood flow (from the lobule periphery to the lobule center; Felbeck and Turner, 1995). This presumptive concentration gradient may lead to differential availability of inorganic carbon (and other substrates, see below), which in turn likely results in differential regulation of bacterial gene expression, such as highest abundance of CO_2_ incorporation enzymes in large symbionts. Large *Riftia* symbionts thus presumably not only benefit from higher CO_2_ levels, but also have more biosynthetic capacities at their disposal than small symbionts: Small Endoriftia need to maintain cell division and, consequently, invest a considerable part of their resources in growth-related processes and the expression of putative host interaction-related proteins that ensure survival of the stem cell population (see above). In contrast, large symbionts apparently divide less frequently and may be less endangered of host interference (before they reach the degenerative lobule zone) and can thus allocate more energy to production of organic material. This would eventually benefit the host, which digests the larger symbionts at the trophosome lobule periphery. Our observation that most cofactor- and vitamin metabolism-related proteins were more abundant in fractions M and/or L than in fractions XS or S supports the idea of relatively more biosynthesis in large Endoriftia.

Higher abundances of glycogen-producing enzymes in fraction L furthermore suggest that large symbionts invest relatively more of their biosynthetic capacities in storage of fixed carbon in the form of glycogen than smaller symbionts. This is in accordance with a previous study (Sorgo *et al.*, 2002), which noted a glycogen gradient in the symbiont cells, with increasing glycogen density from the lobule center towards the periphery, i.e., towards larger symbiont cells.

#### Small Endoriftia store more sulfur and are more involved in sulfide oxidation

Smaller symbionts produce relatively more sulfur globules for sulfur storage than larger symbiont cells, as indicated by relatively higher abundance of sulfur globule proteins in fraction XS (Supplementary Figure S4). This is in agreement with observations of Hand (1987), who noted more sulfur deposits in central (small) than in peripheral (large) *Riftia* symbionts. Although this finding was not supported by a subsequent study (Pflugfelder *et al.*, 2005), our results do point to different amounts of S storage in different Endoriftia subpopulations. As shown for the free-living thiotrophic model bacterium *Allochromatium vinosum*, activation of stored sulfur involves trafficking proteins such as TusA, which is involved in sulfur transfer to DsrEFH and DsrC (Stockdreher *et al.*, 2014). In our study, the highly abundant TusA, several DsrC copies as well as DsrEFH were all detected with highest abundances in fraction XS, thus supporting the idea of relatively more re-mobilization of sulfur and subsequent utilization of reduced sulfur compounds in small Endoriftia. As the highly abundant adenylylsulfate reductase AprAB, the ATP-sulfurylase SopT and sulfide dehydrogenase subunit FccB were also detected with higher abundances in fractions XS or S than in M or L, one might conclude that sulfide oxidation itself also plays a more prominent role in smaller symbionts than in larger symbionts. However, as we detected the dissimilatory sulfite reductase DsrAB, the third key enzyme of cytoplasmic sulfide oxidation, with a rather ambiguous abundance pattern (Supplementary Table S4e), this idea remains speculative and requires further analysis.

#### In large symbionts, thiosulfate oxidation plays a more prominent role

Larger symbionts may rely relatively more on thiosulfate oxidation – in addition to sulfide oxidation – than smaller Endoriftia, as suggested by highest abundance of SoxZ and detection of several other (low-abundant) Sox proteins in fraction L. Expression of the Sox (sulfur oxidation) complex was shown to be upregulated in the presence of thiosulfate in *A. vinosum* (Grimm *et al.*, 2011). We speculate that thiosulfate concentrations might be higher in the trophosome lobule periphery than in the lobule center, due to a concentration gradient (as proposed above for CO_2_) and/or possibly also as a result of host thiosulfate production. The *Riftia* host appears to be able to oxidize toxic sulfide to the less toxic thiosulfate in its mitochondria (Hinzke *et al.*, 2019). Higher abundance of host thiosulfate sulfurtransferase in symbiont-enriched fractions compared to non-enriched trophosome homogenate in our present study suggests that this putative detoxification process could be particularly important in the symbiont-containing bacteriocytes. With sulfide supposedly reaching the trophosome lobule periphery first with the blood flow, free sulfide concentrations might be higher there and, consequently, host sulfide oxidation to thiosulfate might be more frequent in bacteriocytes at the lobule periphery than in the center. The idea of more thiosulfate oxidation in large Endoriftia is further substantiated by highest abundance of six rhodanese family proteins in fraction L, as rhodanese-like proteins can cleave thiosulfate into sulfite and sulfide and were proposed to be involved in thiosulfate oxidation (Hensen *et al.*, 2006, Welte *et al.*, 2009).

Interestingly, overall abundance of all proteins involved in the symbiont’s energy-generating sulfur metabolism, the most abundant of all metabolic categories, remained relatively unchanged across the four fractions (Supplementary Table S7). This indicates that sulfur oxidation-based energy generation, a fundamental basis of all other metabolic processes, is equally important throughout the symbiont’s differentiation process, even if individual contributions of reduced sulfur compounds may differ. (For a detailed overview of sulfur oxidation reactions in Endoriftia see Supplementary Results and Discussion F).

#### *Hydroge*n *oxidation is more relevant in large symbionts*

In large symbionts, the use of hydrogen may furthermore play a more prominent role than in smaller symbiont cells, as suggested by increasing abundances of the Isp-type respiratory H_2_-uptake [NiFe] hydrogenase large subunit HyaB, a Fe-S oxidoreductase (GlpC) encoded next to HyaB, and the hydrogenase expression/formation protein HypE from fraction XS to L. The small hydrogenase subunit HyaA (Sym_EGV51837.1) and an additional hydrogenase expression/formation protein (HoxM, Sym_EGV51835.1), both of which are encoded upstream of HyaB in the symbiont genome, were detected with increasing abundance towards fraction L as well (although at very low concentrations; Supplementary Table S3b), supporting the idea of relatively more hydrogen oxidation in large symbionts. Like for CO_2_ and thiosulfate, this might be due to a concentration gradient with highest hydrogen concentrations at the lobule periphery and lowest concentrations towards the lobule center. Use of hydrogen as an energy source has been described or suggested for free-living sulfur oxidizing bacteria like *A. vinosum* (Weissgerber et al., 2011), and for a variety of thiotrophic symbionts of marine invertebrates (Petersen *et al.*, 2011). Taking advantage of hydrogen oxidation in addition to sulfide- and thiosulfate oxidation, i.e., using a broader repertoire of electron donors, would potentially enhance the metabolic flexibility, particularly of large Endoriftia. However, H_2_ was recently suggested to be involved in maintaining intracellular redox homeostasis rather than working as electron donor in the *Riftia* symbiosis (Mitchell *et al.*, 2019), and hydrogenase may in fact also play a role in sulfur metabolism (as suggested for *A. vinosum* (Weissgerber *et al.*, 2014); Supplementary Results and Discussion F). Therefore, the exact role of hydrogen oxidation in Endoriftia and why it might be relatively more relevant in larger symbionts remains to be discussed.

#### Denitrification in Riftia symbionts appears to be modular

Our results suggest that small *Riftia* symbionts rely relatively more on the NarGHI-mediated first step of respiratory nitrate reduction to nitrite, while all subsequent steps of nitrite reduction via NO and N_2_O to N_2_ seem to be more prominent in larger symbionts. Since expression of nitrate reduction genes is usually inhibited by oxygen (Payne, 1973), high NarGHI abundance in fraction XS suggests that O_2_ levels might be particularly low in the trophosome lobule center.

Interestingly, although Endoriftia has the genomic potential for complete denitrification to N_2_, small and large symbionts seem to employ separate parts of the pathway. This is reminiscent of free-living microbial communities, in which denitrification is modular, i.e., it is often not carried out by individual organisms, but rather by the subsequent activity of several members (Graf *et al.*, 2014), between which intermediates are passed on as ‘metabolic handoffs’ (Anantharaman *et al.*, 2016). Moreover, nitrate reduction and subsequent denitrification steps may occur as two temporally separated processes even in the same organism: During nitrate reduction in *Staphylococcus carnosus*, nitrite reduction was inhibited and resumed only after nitrate was depleted (Neubauer and Götz, 1996). A similar scenario might be assumed for Endoriftia: Small symbionts apparently reduce nitrate to nitrite, which, potentially, yields enough energy to cover their demand, while ‘saving’ nitrite as a handoff for future use. Once the symbionts have become larger, expression of nitrite reductase, nitric oxide reductase and nitrous oxide reductase in higher abundance enables them to further reduce the accumulated intermediate nitrite. Whether these reactions could also be regulated in response to varying oxygen concentrations is unclear. We speculate that an O_2_ gradient may exist, which influences the observed expression pattern.

#### Regulation of gene expression may be less stringent in large symbionts

Relative abundance of the RNA polymerase sigma factor RpoD decreased from fraction XS to L (Figure 3, Supplementary Table S4h) in S-rich and S-depleted samples, pointing to relatively more growth-related activities in small Endoriftia (see also Supplementary Results and Discussion B). RpoD is the primary sigma factor for vegetative growth (σ70), which regulates transcription of most genes involved in exponential growth in many bacteria (Helmann and Chamberlin, 1988, Fujita *et al.*, 1994, Ishihama, 2000). This would be in agreement with the idea of small *Riftia* symbionts being mainly occupied with cell division and proliferation in a quasi-exponential growth phase, while large symbionts function as biosynthetic ‘factories’, focusing on carbon fixation and biomass production. Interestingly, RpoS, the master transcriptional regulator of stationary phase gene expression and antagonist of RpoD, was not detected in any of our samples (although it is encoded in the symbiont genome). RpoS abundance increases upon stress and limitation during transition to the stationary phase in free-living model bacteria (Hengge-Aronis, 1993, Fujita *et al.*, 1994, Ishihama, 2000). Its absence in the *Riftia* symbiont’s proteome suggests that, unlike free-living bacteria, the symbiont does not experience a stationary phase-like growth arrest even in later developmental stages, probably because it is ideally supplied with all necessary substrates by the host. This ‘lack’ of stress or limitation possibly results in less stringent regulation of symbiont gene expression, which could explain the metabolic diversity we observed particularly in large symbionts, such as multiple ways of energy generation (thiosulfate- and hydrogen oxidation in addition to sulfide oxidation) and two CO_2_ fixation pathways. Under these premises, the previously observed simultaneous expression of seemingly redundant metabolic pathways in *Riftia* symbionts (Markert *et al.*, 2011) very likely reflects this presumptive “de-regulation” of gene expression in large parts of the symbiont population, which allows Endoriftia to fully exploit its versatile metabolic repertoire to the advantage of the symbiosis.

## Conclusion

Our results show that Endoriftia cells of different differentiation stages likely employ distinct metabolic profiles, thus confirming our initial hypothesis. Whereas small Endoriftia ensure survival of the symbiont population, large Endoriftia are primarily engaged in biomass production. The driving force behind this differentiation remains to be elucidated. For *Rhizobium*, a steep O_2_ concentration gradient inside legume nodules was proposed to be involved in signaling for symbiont differential gene expression (Soupene *et al.*, 1995). Similarly, some of the differences we observed in small and large Endoriftia might also be connected to the availability of electron donors or acceptors, and hence differentiation of Endoriftia cells might depend on substrate availability. Symbiont differentiation in *Riftia* might furthermore be induced by specific host effectors, e.g., histone-derived antimicrobial peptides, which were recently proposed to play a role in symbiont cell cycle regulation (Hinzke *et al.*, 2019), or other compounds that allow *Riftia* to modulate the symbiont’s expression of certain metabolic pathways. Besides such direct interference, *Riftia* likely also exerts indirect influence on symbiont gene expression by providing copious amounts of all necessary substrates to the bacterial partner. We speculate that this constantly high nutrient availability inside the host causes Endoriftia’s biosynthetic pathways to be regulated less stringently (compared to what we would expect in free-living bacteria). This would explain the previously observed metabolic versatility of symbionts in the same host: Large Endoriftia can afford to employ much of their metabolic repertoire at the same time. Such an ‘advantageous deregulation’, i.e., unhindered expression of multiple – even redundant – metabolic pathways, likely enables high symbiont productivity during symbiosis.

## Supporting information

Supplementary Results and Discussion with Supplementary Figures S1-S4 and Supplementary Tables S1, S2, S6-S9

## Acknowledgements

We thank captains and crews of R/V Atlantis and DSV Alvin who supported sampling during cruises AT26-23 and AT37-12. We are grateful to Jana Matulla and Annette Meuche for excellent technical assistance in sample preparation for proteomics and electron microscopy, respectively. Thanks to Ruby Ponnudurai and Frank Unfried for help with CARD-FISH, and to Alexander Graf, Mathis Appelbaum, Judith Zimmermann and Silke Wetzel for advice on epifluorescence microscopy and staining. We greatly appreciate Elisa Kasbohm’s help with random forest analyses. We are very grateful to Jörg Bernhardt for stitching the transmission electron micrographs to produce a panorama image with high resolution. This work was supported by the German Research Foundation DFG (grant MA 6346/2-1 to S.M.), a fellowship of the Institute of Marine Biotechnology Greifswald (T.H., M.M.), a German Academic Exchange Service (DAAD) grant (T.H.), the NC State Chancellor’s Faculty Excellence Program Cluster on Microbiomes and Complex Microbial Communities (M.K.), the USDA National Institute of Food and Agriculture, Hatch project 1014212 (M.K.), the U.S. National Science Foundation (grants OCE-1131095 and OCE-1559198 to S.M.S), and The WHOI Investment in Science Fund (to S.M.S).

## Author contributions

S.M. conceived the study, S.M., T.H., M.K. designed the experiments, T.H. and H.F. took samples. T.H. prepared samples for CARD-FISH and metaproteomics analysis, analyzed data, conducted statistical analyses and prepared figures. T.H. and S.M. wrote the manuscript with input from all coauthors. T.S. was involved in project coordination, S.M.S. obtained funding for the research cruises and coordinated sampling as chief scientist. C.H. and F.B. performed MS measurements of metaproteomics samples, D.B. coordinated MS measurements. M.M. and J.P.-F. contributed to fluorescence microscopy and R.S. conducted electron microscopy analyses. P.H. performed flow cytometry analyses, and U.V. coordinated flow cytometry measurements.

