## Supplementary Results and Discussion with Supplementary Figures S1-S4 and Supplementary Tables S1, S2, S6-S9 for "Metabolic differences between symbiont subpopulations in the deep-sea tubeworm *Riftia pachyptila*"

### Contents

This file:

Supplementary Results and Discussion with Supplementary Figures S1 – S4

Supplementary Table S1: Sampling details

Supplementary Table S2: Nucleotide sequences of HCR-FISH probes

Supplementary Table S6: Total numbers of identified proteins

Supplementary Table S7: Summed up abundances of metabolic categories

Supplementary Table S8: Flow cytometry data

Supplementary Table S9: Sulfur metabolism-related Endoriftia proteins

The following Supplementary Figure is provided as separate PDF file:

Supplementary Figure S5: High resolution version of TEM image (Figure 5a)

The following Supplementary Data Sets are provided as separate Excel files:

Supplementary Table S3: All detected Endoriftia proteins with relative abundances

Supplementary Table S4: Abundance trends of symbiont proteins in the categories
(a) cell cycle & DNA metabolism
(b) chaperones
(c) transport
(d) carbon metabolism
(e) chemotrophy
(f) nitrogen metabolism
(g) cofactors and vitamins
(h) transcription & translation

Supplementary Table S5: *Riftia* host proteins with higher abundances in XS and S

### Supplementary Results and Discussion

#### A) Symbiotic Endoriftia cells exist in a remarkable size range

Symbiont cell sizes in *Riftia* trophosome tissue range from 1-2 µm to more than 15 µm (main text: Figure 1, Figure 5). This is in line with previous microscopy-based observations, which suggested that the symbiont cells differentiate from small rod-shaped cells in the trophosome lobule center to larger coccoid cells towards the lobule periphery (Bright and Sorgo, 2003). With an about 10-fold increase in diameter, Endoriftia cells enlarge their volume by a factor of ~1,000 during their differentiation from smallest to largest coccoid symbiont cells. Considerable enlargement of bacterial cells in the course of symbiotic differentiation has also been observed in the intracellular thiotrophic symbiont of the shallow water clam *Codakia orbicularis* (increases 10-fold in length; Caro *et al.*, 2007), in *Sinorhizobium meliloti* in alfalfa nodules (increases four- to seven-fold in length; Oke and Long, 1999), in symbionts of the nematode *Eubostrichus* (increase up to 13-fold in length; Pende *et al.*, 2014), and in the giant bacterium *Epulopiscium fishelsoni*, intestinal symbiont of surgeon fish (increases up to 3,000-fold in volume; Bresler and Fishelson, 2003). Such enormous size gradients are rather the exception than the rule in bacteria, however. Cell sizes, that is, length or diameter, of free-living model bacteria like *B. subtilis* or *E. coli* usually vary only by factor 2 (during cell division), i.e., these bacteria may increase their volume 2-fold (assuming a cylindrical shape) to 8-fold (assuming a spherical shape) at most (Chien *et al.*, 2012). This suggests that the remarkably large size range observed for Endoriftia presents a consequence of its symbiotic life style.

#### B) Comparative analysis of enriched symbiont fractions from S-rich vs. S-depleted *Riftia* specimens

##### *Overview*

Our comparative analyses of symbiont-enriched fractions XS to L revealed that in both, S-rich and S-depleted samples, protein profiles differed with increasing symbiont cell size (Supplementary Figure S1). Many groups of proteins (e.g. carbohydrate metabolism-related proteins) showed similar trends across size classes in S-rich and S-depleted specimens, even if individual protein abundances differed. Statistical testing for significant differences in protein abundance between S-rich and S-depleted fractions of the same size class returned only very few (edgeR) or no (random forest) hits. This may in part be due to the less effective enrichment of symbionts from S-depleted trophosome tissue homogenate. However, very similar abundance patterns in symbionts from sulfur-rich and sulfur-depleted hosts might also reflect the fact that symbionts are very well buffered against environmental changes (as previously suggested, Hinzke *et al.*, 2019) and, therefore, functional differences between symbiont morphotypes in S-rich vs. S-depleted symbionts might be negligible. Some of these differences, however, seemed to be specific for the respective energy situation and are outlined below.

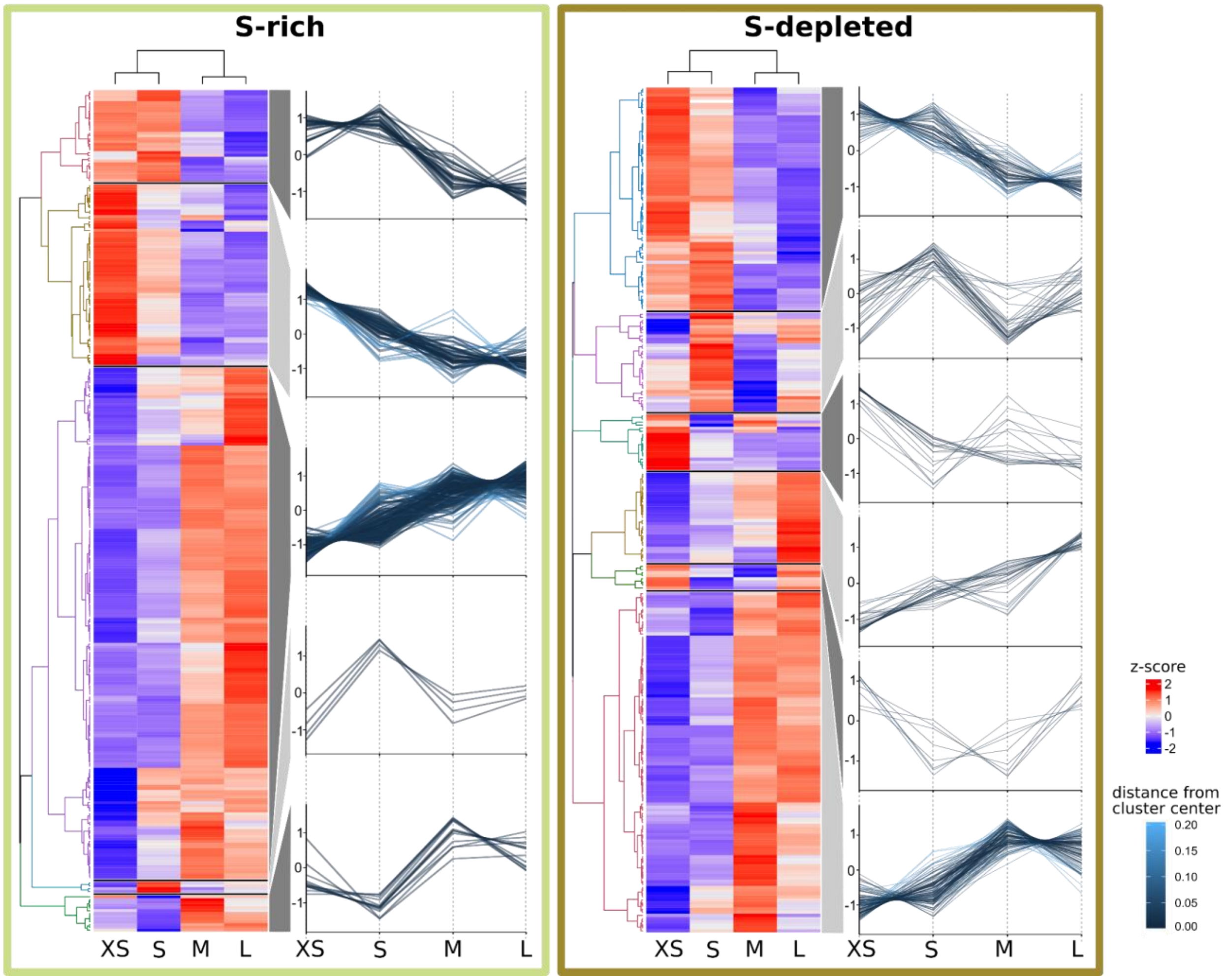

**Supplementary Figure S1:** Abundance trends of 465 *Riftia* symbiont proteins with significant abundance differences between the four analyzed gradient fractions XS (enriched in very small symbiont cells) to L (containing the highest percentage of large symbiont cells) in S-rich and S-depleted *Riftia* trophosomes. Heat maps show relative protein abundances (z-scores of edgeR-RLE-corrected spectral count values; see Methods for details) and line graphs indicate trends in the observed differences.

##### *Cell division*

In sulfur-depleted hosts, *Riftia* symbionts appear to divide less frequently than in sulfur-rich specimens, as indicated by lower abundance of the major cell division protein FtsZ in all S-depleted fractions compared to their S-rich counterparts (Supplementary Table S3; please note that, due to its low abundance, FtsZ was not included in statistical analysis in S-depleted samples). In S-depleted fraction XS, FtsZ abundance was about 3.5 times lower than in S-rich fraction XS. Less symbiont cell division in S-depleted *Riftia* accords with the idea of severe energy limitation in sulfur-depleted symbionts and is in agreement with our previous finding that symbiont proteinaceous biomass is lower in trophosomes of S-depleted specimens (Hinzke *et al.*, 2019). In this previous study, we suggested that S-depleted hosts digest a larger part of their symbiont population as compared to S-rich tube worms. As the host mainly digests large symbionts at the trophosome lobule periphery (Figure 5 main text; Bright and Sorgo, 2003), one might expect that more digestion leads to relatively more smaller symbionts in S-depleted trophosomes as compared to S-rich hosts. This was, however, not the case, as symbiont size distribution was quite comparable in trophosome homogenates of S-rich and S-depleted trophosomes (see Figure 1, main text). We therefore presume that both, more symbiont digestion and less symbiont cell division co-occur in S-depleted worm specimens, leading to the previously observed loss in total symbiont biomass.

##### *Growth-related processes*

Highest abundance of RNA polymerase subunits, transcription elongation factors, transcription antitermination protein and various translation-related proteins in fraction XS of S-rich and S-depleted specimens indicates that small symbionts devote relatively more energy and resources to protein synthesis than large symbionts (Supplementary Table S4h). This is in agreement with the idea that small Endoriftia function as actively dividing and growing stem cells of the Endoriftia population, whereas large symbionts have the role of highly efficient biomass producers (see main text). This proposed greater importance of growth-related processes in small symbionts may result in higher intracellular pyrophoshate levels, as suggested by high abundance of pyrophosphatases in fraction XS. The highly abundant pyrophosphate-energized proton pump HppA (Sym_EGV49909.1) and the inorganic pyrophosphatase Ppa (Sym_EGV49908.1) had their highest abundances in fraction XS in S-rich samples (Supplementary Table S3). Pyrophosphatases play an important role in energy metabolism by catalyzing the hydrolysis of inorganic pyrophosphates ($PP_i$), which are produced at particularly high rates by biosynthetic reactions in growing cells (Klemme, 1976, Chen *et al.*, 1990). By removing $PP_i$, pyrophosphatases shift the thermodynamic equilibrium to favor reactions like DNA, RNA and protein synthesis (Lahti, 1983). HppA may furthermore have an additional growth-related function: During $PP_i$ hydrolysis, HppA pumps protons into the periplasm, thus establishing a proton motive force (Maeshima, 2000). As cell division is an energy-expensive process, which requires not only ATP but also proton motive force (Goehring and Beckwith, 2005), HppA may be upregulated to accommodate this increased demand in small, dividing Endoriftia. At the same time, HppA presumably increases energy efficiency of the Calvin cycle (Markert *et al.*, 2011). Interestingly,

HppA abundance was notably lower in S-depleted XS fractions, supporting the idea of reduced cell division in energy-depleted symbionts (see above).

Besides HppA, highly abundant nitrate reductase NarGHI, which also produces a proton gradient, may provide a similar advantage in small symbionts (see main text).

##### *Host interactions*

Proteins which may protect the symbiont from digestion by the host could be most important in small symbiont cells and particularly so in S-depleted *Riftia*, as suggested by highest abundances of an ankyrin protein and of the FK506-binding protein FkpA in S-depleted fraction S (Supplementary Table S3). In S-rich and S-depleted samples, the ankyrin-like symbiont protein (Sym_EGV51005.1) decreased in abundance from fraction S to L. Endoriftia ankyrin repeat-containing proteins were previously suggested to be involved in microbe-host interactions, possibly to counteract digestion by the host (Hinzke *et al.*, 2019). As small Endoriftia are the main dividing symbiont subpopulation and thus ensure survival of the symbiont population as a whole, digesting those cells would harm not only the symbiont, but also the host itself. The ankyrin protein could fulfill a protective role especially for these smaller symbionts. The *Riftia* symbiont's FK506-binding protein (Sym_EGV50540.1), which showed a comparable abundance trend, might have a similar role. In *Salmonella typhimurium* and *Cronobacter*, FkpA is involved in survival inside host cells (Horne *et al.*, 1997, Eshwar *et al.*, 2015), suggesting that the Endoriftia FkpA, too, provides protection for the intracellular symbiont.

#### C) Flow cytometry of *Riftia* symbionts

According to our flow cytometry data, *Riftia* trophosome homogenate and enriched gradient fractions were quite heterogeneous (Supplementary Figure S2), with a number of other populations present besides population 1 (small symbionts) and 2 (large symbionts). This heterogeneity is presumably due to the fact that a) symbionts exist not only as small or large cells, but also adopt any intermediate size, and b) intracellularly stored sulfur influences the cells' light-scattering properties (especially side scatter, SSC), considerably (as shown for thiotrophic lucinid symbionts; Caro *et al.*, 2007). We sorted one of the additional populations, with SSC between $10^4$ and $10^5$ and FSC between $10^3$ and $10^4$ (i.e. with higher SSC but lower FSC than population 1 and 2) to examine it separately. Fluorescence microscopy revealed that this population consisted mostly of medium-sized symbionts, which – unlike population 1 and 2 - contained numerous sulfur globules (images not shown). It can be assumed that other symbiont cell populations, e.g. small sulfur-rich and large sulfur-rich cells, might also be present. This hypothesis awaits confirmation in future studies. To estimate symbiont DNA content in the present study, we only included populations 1 and 2, which were readily comparably due to their similar sulfur content (i.e., there were hardly any sulfur globules visible).

As also described for a thiotrophic lucinid symbiont (Caro *et al.*, 2007), cell populations were not entirely congruent across the two bioreplicates in our Endoriftia flow cytometry analyses. Consequently, individual fluorescence intensity (FI) values varied considerably (Supplementary Table S8). Nevertheless, both replicates clearly showed the same trend, i.e., higher FI per particle in population 2 compared to population 1 across all samples, strongly indicating multiple genome copy numbers in large symbionts.

**Supplementary Figure S2 (next page):** Fluorescence microscopy and fluorescence-activated cell sorting (FACS) of *Riftia* symbiont cells. Left: Micrographs (Phase contrast and Syto9 staining), right: Flow cytometry dot plot (FSC: forward scatter, SSC: side scatter) and histogram. To identify symbiont subpopulations of different cell sizes, a set of six individual gradient fractions enriched in large symbiont cells and a set of six gradient fractions enriched in small symbiont cells were examined and compared (note that only one pair of microscopy images and only one of the six dot plot/histogram pairs per sample set are shown.) **A)** Those fractions that were enriched in small symbiont cells of 2-3 µm in diameter produced dot plots with a highly abundant cell population 1 (encircled in black), which we assumed to be specific for small symbionts. **B)** In contrast, gradient fractions enriched in large symbionts of up to 10 µm in diameter produced dot plots in which population 1 was notably less prominent, while a second population (2) was highly abundant. Population 2 was almost completely absent in (A) and therefore presumably specific for large symbionts. **C)** Non-enriched trophosome homogenate contained a mixture of cells and particles of different sizes. Both cell populations determined in (A) and (B), presumably indicative of small (1) and large (2) symbionts, were also visible in the homogenate's dot plot and histogram, which allowed us to measure and compare their respective fluorescence signal intensities. Median fluorescence intensity per particle of population 1 was consistently (throughout all samples) lower than that of population 2, even if cell counts for population 1 were higher than for population 2 (e.g., A, D). The green fluorescent dye Syto9 stains DNA and RNA, and thus – since RNA had been removed from the samples by RNase treatment before analysis – enabled us to quantify DNA content in populations 1 and 2 (see Supplementary Table S8). **D** and **E)** Sorting of the two populations 1 and 2 from trophosome homogenate and subsequent examination of the resulting sorted cell suspensions by microscopy and flow cytometry confirmed that these two populations are indeed small (D) and large (E) symbiont cells. Trophosome homogenate and gradient fractions used in this analysis originated from two *Riftia* specimens with medium sulfur content (see Supplementary Table S1 for details). Scale bar: 10 µm.

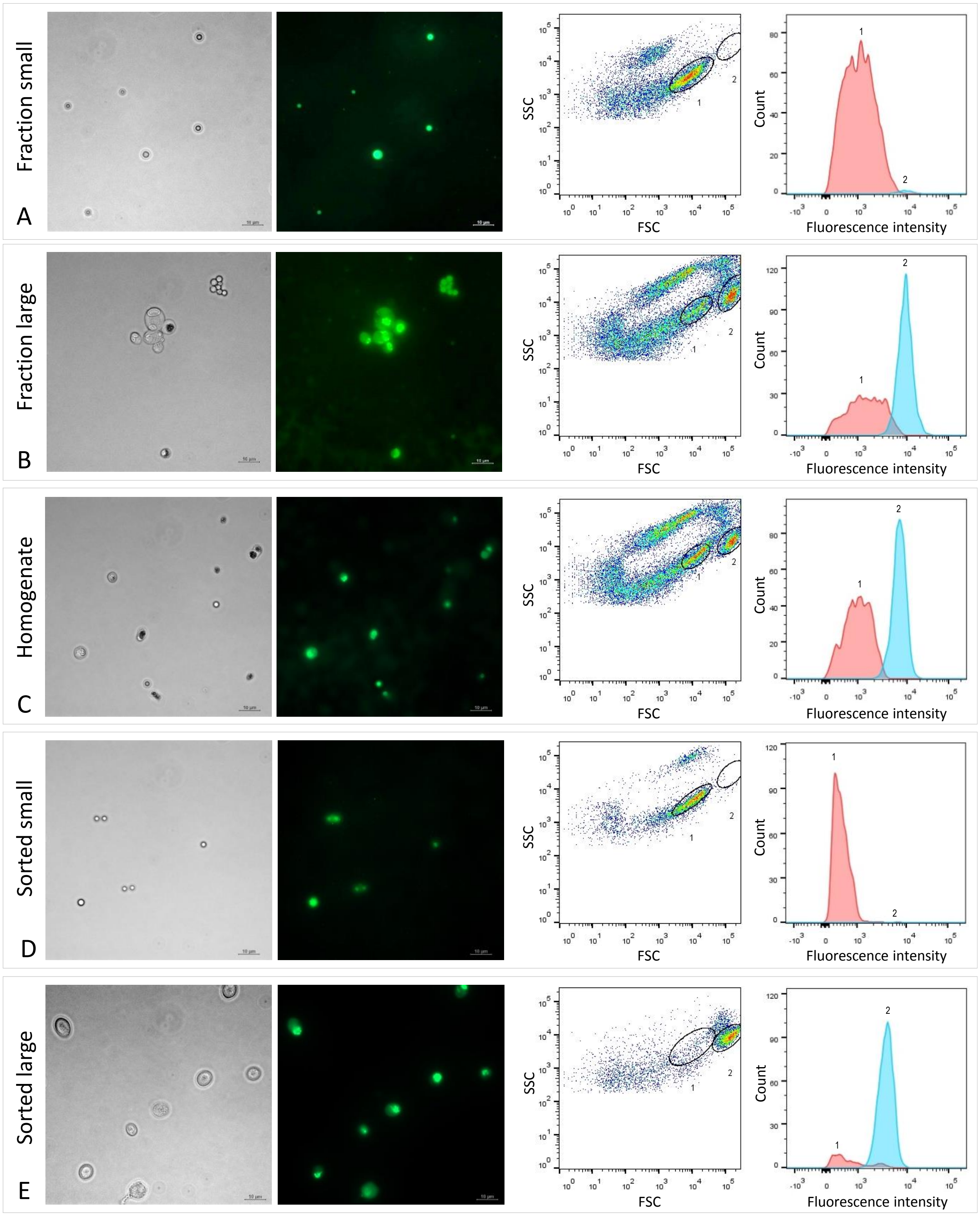
Fraction small
A
Fraction large
B
Homogenate
C
Sorted small
D
Sorted large
E
SSC
FSC
Count
Fluorescence intensity

#### D) $CO_2$ metabolism is differentially regulated across Endoriftia cell sizes

##### *The carbon fixation key enzyme RubisCO is more abundant in large Endoriftia*

The Calvin cycle key enzyme RubisCO was detected with notably higher mRNA-based fluorescence intensities in large Endoriftia cells, compared to smaller symbionts. This is in agreement with our proteomic results (see main text), and supports the conclusion that large symbionts are more involved in carbon fixation and, generally, in biomass production, than small symbionts (Supplementary Figure S3).

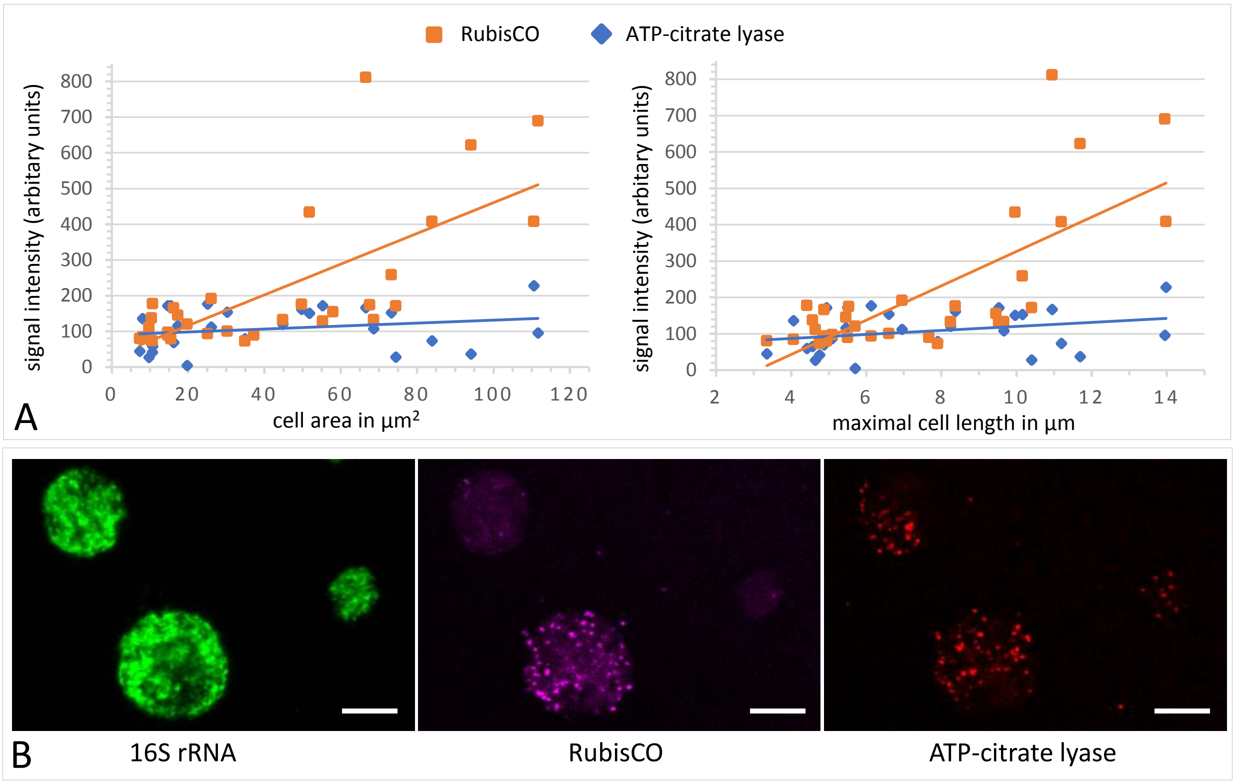

**Supplementary Figure S3**: A gradient fraction enriched in large symbionts (but also containing small symbiont cells) was fixed as for CARD-FISH analysis and incubated with fluorescently labelled RNA probes against the Endoriftia 16S rRNA and the mRNAs of Calvin cycle key enzyme RubisCO and rTCA cycle key enzyme ATP-citrate lyase (subunit AclB) before examination by confocal laser scanning microscopy (CLSM, see Methods). **A)** Background-corrected mean signal intensities per pixel calculated from a total of 33 cells (in eight images) plotted against cell area (left) and Feret's Diameter of the cell (right). Straight lines indicate the linear between mean pixel intensities and cell size. Average RubisCO mRNA signal intensity increased notably with cell size (orange lines), while AclB signal intensity increased only very slightly (blue lines). **B)** CLSM image of Endoriftia cells. Supporting the quantitation in A) and in line with our proteomic results, the RubisCO signal is markedly more intense in large symbiont cells than in small cells, while the AclB signal is very weak and signal intensity differences between large and small cells seem to be minor. Scale bar = 5 µm. Image brightness and contrast were manually adjusted.

*Expression patterns of TCA cycle enzymes are ambiguous*

Like RubisCO, the rTCA cycle key enzyme ATP-citrate lyase small subunit (AclA, Sym_2601634392) was detected with significantly increasing abundance from fraction XS to L in our proteomic analyses, suggesting that carbon fixation plays a relatively more important role in large *Riftia* symbionts than in small symbionts (see main text). However, expression of other (r)TCA cycle enzymes was surprisingly inconstistent, i.e., while abundance of some enzymes increased towards fraction L (including the key enzymes AclA and KorAB), other enzymes showed the opposite – albeit non-significant – trend and became less abundant (e.g., AclB, isocitrate dehydrogenase Icd; Supplementary Table S4d, Supplementary Table S3). Further contributing to this ambiguous pattern, the AclB mRNA signal was detected with very similar (and very low) abundances in small and large Endoriftia cells (see Supplementary Figure S3 above). A possible explanation for these observations might be that Endoriftia's (r)TCA cycle enzymes can run in either direction, depending on cellular requirements. While certain key reactions of TCA and rTCA cycle have long been considered as irreversible, this seems not always to be the case, as, for instance, reported for citrate synthase, key enzyme of the oxidative TCA cycle, which can also operate in the reverse direction, cleaving citrate (Mall *et al.*, 2018). Endoriftia's citrate synthase (although encoded in the genome) was not detected at all on the protein level in this study, allowing for the speculation that AclAB might functionally replace citrate synthase in the oxidative version of the TCA cycle by running in reverse, possibly even producing ATP in the process. Assuming that the observed discrepancies in Endoriftia (r)TCA cycle enzyme abundance trends are thus indeed caused by flexible changes in the enzymes' operating directions, Icd could, for example, produce oxaloacetate (e.g., for glutamate synthesis) and NADH in small symbionts, while in large symbionts, Icd might fix $CO_2$ by running in the reverse direction. Further studies are required to solve the exact regulation of the symbiont (r)TCA cycle. The recently described combination of matrix-assisted laser desorption/ionization mass spectrometry and FISH (metaFISH), which allows for discrimination of symbiont subpopulations based on the metabolites they produce (Geier *et al.*, 2020), might be a promising tool for this purpose.

*Large symbionts may take up organic compounds in addition to $CO_2$*

Our detection of five *Riftia* symbiont TRAP transporter subunits and four ABC transporter components putatively involved in uptake of organic material with increasing relative abundance from fraction XS to L indicates that Endoriftia imports small organic compounds, particularly in the late stage of differentiation, i.e., in large cells. ABC transporters can mediate uptake of small molecules (such as sugars, amino acids or vitamins), and metal ions (Davidson *et al.*, 2008), while TRAP transporters facilitate import of C4-dicarboxylates like fumarate, succinate and malate (Dct type; Mulligan *et al.*, 2011) or of amino acids like glutamate and glutamine (TAXI type; Mulligan *et al.*, 2007). All of these compounds may be relevant heterotrophic substrates in large Endoriftia, which could channel amino acids and peptides into protein biosynthesis, while sugars could be stored as glycogen. Heterotrophy in thiotrophic symbionts was previously shown for a ciliate symbiont (Seah *et al.*, 2019) and for ectosymbionts of shrimp (Ponsard *et al.*, 2013). Although the *Riftia* symbiont's potential for mixotrophy, i.e., for both, autotrophy and heterotrophy, had been predicted from the symbiont's genome, it was previously assumed that heterotrophy might be particularly relevant in free-living Endoriftia, but not during symbiosis (Robidart *et al.*, 2008). Our results challenge this assumption and suggest that Endoriftia relies on mixotrophy even when in symbiosis, which would allow re-cycling of carbon from host to symbiont.

#### E) Small Endoriftia might be nitrogen-limited

Small Endoriftia may rely relatively more on the glutamine synthetase-glutamate synthase (GS-GOGAT) pathway for ammonia assimilation, while large symbionts cells seem to preferably use glutamate dehydrogenase (GDH) for this purpose. Both, in S-rich and in S-depleted samples, a glutamine synthetase copy (GlnA), glutamate synthase subunit GltB and nitrogen regulatory protein P-II (GlnB) were detected with decreasing abundance from fraction XS to L (Figure 3 main text, Supplementary Table S4f). In contrast, glutamate dehydrogenase (GdhA) showed the opposite trend with lowest abundance in XS and highest abundance in L (S-rich) or M (S-depleted). The GS-GOGAT pathway, which is energetically more expensive than GDH, was shown to be used under energy-rich conditions or during nitrogen limitation in *E. coli* (reviewed in Reitzer, 2003). GS-GOGAT was furthermore shown to have a higher affinity towards ammonium than GDH (Reitzer, 2003). This suggests that small symbionts could be nitrogen-limited, either due to a concentration gradient (with highest nitrogen levels in the peripheral lobule zones), and/or due to their own high demand for nitrogen compounds for growth. Further investigations are required to evaluate this speculation.

#### F) Sulfur metabolism

While many of the energy-generating reactions of the uncultured *Riftia* symbiont's sulfur metabolism have been elucidated previously (Markert *et al.*, 2011), several details remained vague. Our new proteome data enabled us to propose a more detailed model of the Endoriftia sulfur metabolism (Supplementary Figure S4, Supplementary Table S9).

*DsrC:* The Endoriftia genome encodes several copies of DsrC family proteins, four of which were detected as proteins in this study (Supplementary Table S3). One of them, Sym_EGV52266.1, was one of the most abundant symbiont proteins, pointing to considerable physiological importance of this protein. Similar to the situation in Endoriftia, three putative DsrC copies were found in the *C. okutanii* symbiont (Harada *et al.*, 2009), and DsrC was also the single most abundant sulfur metabolism mRNA in the *Solemya velum* symbiont (Stewart *et al.*, 2011). DsrC has been described to fulfill a key role in dissimilatory sulfur metabolism, including a putative function in transcription regulation and a function as a sulfur trap to allow for maximum DsrAB efficiency (Venceslau *et al.*, 2014). Considering this role of DsrC as enhancer of sulfide oxidation efficiency, highest abundance of all Endoriftia DsrC copies in fraction XS (and lowest DsrC abundance in fraction M or L), corroborates our hypothesis of relatively more $H_2S$ oxidation for energy generation in small *Riftia* symbionts (see main text).

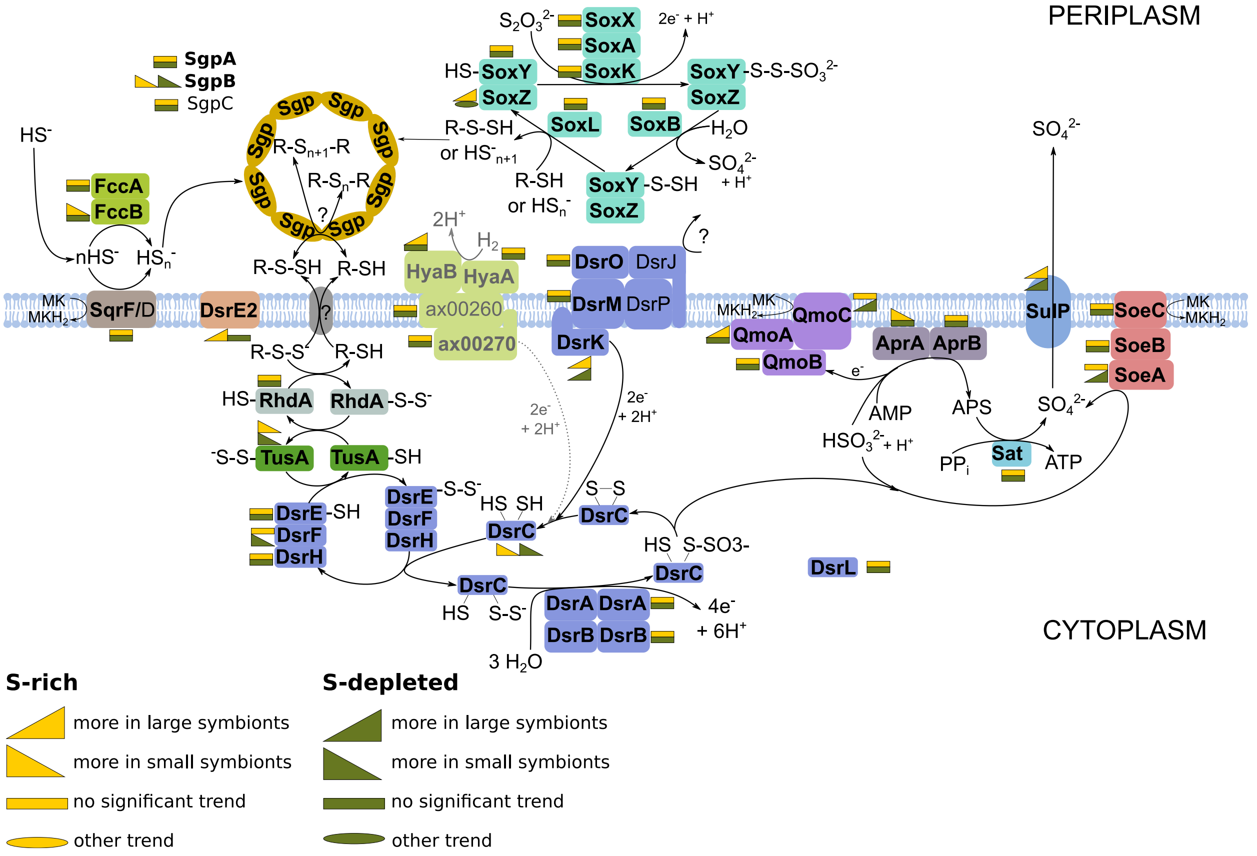

**Supplementary Figure S4**: Energy-generating oxidation of reduced sulfur compounds in Endoriftia. Proteins in bold were detected in this study. (Figure adapted from Grein *et al.*, 2010, Markert *et al.*, 2011, Rodriguez *et al.*, 2011, Stewart *et al.*, 2011, Dahl *et al.*, 2013, Stockdreher *et al.*, 2014, Weissgerber *et al.*, 2014). As the role of hydrogen as electron donor in the *Riftia* symbioses was recently questioned (Mitchell *et al.*, 2019), the associated reactions are labeled in grey.

*SoeABC:* In addition to AprAB and SopT, two of the key enzymes of cytoplasmic sulfide oxidation, we also found SoeABC to be expressed in Endoriftia. In *Allochromatium vinosum*, SoeABC catalyzes direct oxidation of sulfite to sulfur, independently of AMP (Dahl *et al.*, 2013).

*SreABC:* We found the putatively sulfur oxidation-related proteins SreABC in the metagenome and detected SreA on the protein level in Endoriftia. While the exact function of SreABC in the oxidation of reduced sulfur compounds is unclear, for *A. vinosum* it was speculated that the Sre proteins could oxidize polysulfides, which are intermediates generated during sulfide oxidation to sulfur (Weissgerber *et al.*, 2013).

*HyaAB:* Endoriftia's uptake hydrogenase HyaAB might be involved in sulfur oxidation. In *A. vinosum*, concentration of the Isp-type hydrogenase HydLS was shown to increase substantially in the presence of sulfide (Weissgerber *et al.*, 2014), leading to the proposition that hydrogen-derived electrons may be fed into sulfide oxidation via hydrogenase as illustrated in Supplementary Figure S4. *A. vinosum's* HydL (Alvin_2036) and Endoriftia's HyaB (EGV51840.1) protein sequences are 75.69% identical (NCBI BlastP), indicating that both may have similar functions in sulfur oxidation.

**Supplementary Table S1:** Sampling details for specimens and sample types used in this study. All animals were collected at the Crap Spa vent site in the East Pacific Rise (EPR) Tica area. For proteomic analyses, *Riftia* trophosome homogenate (Hom) was subjected to Histodenz-based density gradient centrifugation, separating symbiont cells according to their sizes. After centrifugation, the gradient was carefully disassembled into 24 subsamples/fractions (numbered 1 to 24), all of which were analyzed by CARD-FISH to identify those fractions in which the percentage of very small, small, medium-sized and large symbionts cells was highest. These fractions were designated XS, S, M and L for the respective worm and included in comparative proteomic analyses.

| Analysis | Sampling time | Atlantis cruise number | Trophosome sulfur content | Worm # (biological replicate) | Fraction | Sample descrition/quantile | Dive number (date) | Sampling site | Water depth (m) | Latitude | Longitude |
|---|---|---|---|---|---|---|---|---|---|---|---|
| Proteomics of rate-zonal density gradient fractions | 11/2014 | AT26-23 | S-rich | R#18 (replicate 1) | Hom<br>RZ07<br>RZ08<br>RZ20<br>RZ23 | Homogenate<br>XS<br>S<br>M<br>L | 4764 (09.11.2014) | Crab Spa | 2505 | 09-50.398N | 104-17.479W |
| | | | | R#27 (replicate 2) | Hom<br>RZ08<br>RZ10<br>RZ24<br>RZ22 | Homogenate<br>XS<br>S<br>M<br>L | 4766 (11.11.2014) | Crab Spa | 2512 | 09-50.309N | 104-17.527W |
| | | | | R#28 (replicate 3) | Hom<br>RZ07<br>RZ10<br>RZ24<br>RZ16 | Homogenate<br>XS<br>S<br>M<br>L | 4768 (13.11.2014) | Crab Spa | 2512 | 09-50.373N | 104-17.490W |
| | | | | R#29 (replicate 4) | Hom<br>RZ07<br>RZ09<br>RZ19<br>RZ17 | Homogenate<br>XS<br>S<br>M<br>L | 4768 (13.11.2014) | Crab Spa | 2512 | 09-50.373N | 104-17.490W |
| | | | S-depleted | R#30 (relicate 1) | Hom<br>RZ12<br>RZ15<br>RZ23<br>RZ24 | Homogenate<br>XS<br>S<br>M<br>L | 4769 (14.11.2014) | Crab Spa | 2513 | 09-50.588N | 104-17.434W |
| | | | | R#31 (replicate 2) | Hom<br>RZ07<br>RZ09<br>RZ14<br>RZ17 | Homogenate<br>XS<br>S<br>M<br>L | 4769 (14.11.2014) | Crab Spa | 2513 | 09-50.588N | 104-17.434W |
| | | | | R#32 (replicate 3) | Hom<br>RZ08<br>RZ09<br>RZ24<br>RZ18 | Homogenate<br>XS<br>S<br>M<br>L | 4772 (18.11.2014) | Crab Spa | 2507 | 09-50.449N | 104-17.543W |
| Catalyzed reporter deposition fluorescence in situ hybridization (CARD-FISH) | 11/2014 | AT26-23 | S-rich | R#18 | Hom<br>RZ07<br>RZ08 | Homogenate | 4764 (09.11.2014) | Crab Spa | 2505 | 09-50.398N | 104-17.479W |
| | | | | R#27 | RZ09<br>RZ10<br>RZ11 | | 4766 (11.11.2014) | Crab Spa | 2512 | 09-50.309N | 104-17.527W |
| | | | | R#28 | RZ12<br>RZ13<br>RZ14 | Gradient fractions enriched in different cell sizes from smaller symbiont cells (top) to larger symbiont cells (bottom)<br><br>Hom and all 24 fractions were analyzed in 5 biological replicates. | 4768 (13.11.2014) | Crab Spa | 2512 | 09-50.373N | 104-17.490W |
| | | | | R#29 | RZ15<br>RZ16<br>RZ17 | | 4768 (13.11.2014) | Crab Spa | 2512 | 09-50.373N | 104-17.490W |
| | | | | R#35<br><br>(replicates 1-5) | RZ18<br>RZ19<br>RZ20<br>RZ21<br>RZ22<br>RZ23<br>RZ24 | | 4773 (19.11.2014) | Crab Spa | 2504 | 09-50.449N | 104-17.544W |
| | | | S-depleted | R#30 | Hom<br>RZ07<br>RZ08 | Homogenate | 4769 (14.11.2014) | Crab Spa | 2513 | 09-50.588N | 104-17.434W |
| | | | | R#31 | RZ09<br>RZ10<br>RZ11 | | 4769 (14.11.2014) | Crab Spa | 2513 | 09-50.588N | 104-17.434W |
| | | | | R#32<br><br>(replicates 1-3) | RZ12<br>RZ13<br>RZ14<br>RZ15<br>RZ16<br>RZ17<br>RZ18<br>RZ19<br>RZ20<br>RZ21<br>RZ22<br>RZ23<br>RZ24 | Gradient fractions enriched in different cell sizes from smaller symbiont cells (top) to larger symbiont cells (bottom)<br><br>Hom and all 24 fractions were analyzed in 3 biological replicates. | 4772 (18.11.2014) | Crab Spa | 2507 | 09-50.449N | 104-17.543W |
| Flow cytometry | 11/2014 | AT26-23 | medium S | R#19 (replicate 1) | Hom<br>RZ07<br>RZ08<br>RZ09<br>RZ19<br>RZ21<br>RZ22 | Homogenate<br>Gradient fractions enriched in smaller symbionts (RZ07–RZ09)<br>Gradient fractions enriched in larger symbionts (RZ19–RZ22) | 4764 (09.11.2014) | Crab Spa | 2505 | 09-50.398N | 104-17.479W |
| | | | | R#21 (replicate 2) | Hom<br>RZ07<br>RZ08<br>RZ09<br>RZ19<br>RZ20<br>RZ22 | Homogenate<br>Gradient fractions enriched in smaller symbionts (RZ07–RZ09)<br>Gradient fractions enriched in larger symbionts (RZ19–RZ22) | 4765 (10.11.2014) | Crab Spa | 2511 | 09-50.398N | 104-17.480W |
| Hybridization chain reaction (HCR) FISH | 11/2014 | AT26-23 | S-rich | R#35 | RZ22 | Gradient fraction enriched in larger symbionts | 4773 (19.11.2014) | Crab Spa | 2504 | 09-50.449N | 104-17.544W |
| Transmission electron microscopy (TEM) | 04/2017 | AT37-12 | S-rich | R#9 | - | Intact trophosome tissue | 4895 (28.04.2019) | Crab Spa | 2520 | 09-50.430N | 104-17.502W |
| | | | S-depleted | R#1 | - | Intact trophosome tissue | 4893 (26.04.2019) | Crab Spa | 2508 | 09-50.239N | 104-17.444W |

**Supplementary Table S2:** Nucleotide sequences used for Hybridization chain reaction (HCR) FISH analyses in this study. See Methods for details.

| | |
|---|---|
| > Endoriftia_RubisCO-1-3 | CAACGGGGTAGGCGATCTTCATCAGCTCTTTGGCTTCATCGATCTCATAG |
| > Endoriftia_RubisCO-1-6 | CACATATCCTGGATGTTGACCGCAGGACCGTCGTACAGGCGCAGGTATTT |
| > Endoriftia_RubisCO-1-11 | ATGCACCCAGCAGACGGGTCATCTTGATGTGTACGAAAGCGGTGTAACCA |
| > Endoriftia_RubisCO-1-14 | AAAGGACTCGAAGGCGCGAGCGAACTCTTTGTGCTCTTTCGCGTACTCGA |
| > Endoriftia_AclB-1-1 | AGACGGCGGTAGATAGACCACCGCCACATTGAATTCACAGCCGTCGTCAA |
| > Endoriftia_AclB-1-6 | CGAACTTCTCCATGAACCACTCTTCCTTGGCGACGGCATTGTCACCGGAA |
| > Endoriftia_AclB-1-8 | CTGGTGTCGGTCGGATCTTCGATGCCTGCTTTCTTGAACAGCTCCATCAT |
| > Endoriftia_AclB-1-12 | GTGGGCGAAACCGGTATGGGTGAGGAAACCGATATAGCCTTTGTTGACCT |
| > Endoriftia_AclB-1-18 | AAACAGGAACGTGGTGAAGGCGGCAGATTCCATGGTCGCGTCGCTGATCT |
| > Endoriftia_16srRNA-1 | TATTAGCTCGGATTTCTCCGAGTTGTCCCCCACTACTGGGCAGATTCCTA |
| > Endoriftia_16srRNA-5 | ACGGAGTTAGCCGGTGCTTCTTCTAAAGGTAACGTCAAGACCCAAGGGTA |
| > Endoriftia_16srRNA-9 | TTTACGGCGTGGACTACCAGGGTATCTAATCCTGTTTGCTACCCACGCTT |
| > Endoriftia_16srRNA-13 | TCGGCTCCCGAAGGCACCAATCTATCTCTAGAAAGTTCCGAGGATGTCAA |
| > Endoriftia_16srRNA-14 | GTTCCCCTAGGGCTACCTTGTTACGACTTCACCCCAGTCATGAATCACAA |

**Supplementary Table S6:** Overview of symbiont protein identification numbers in all sample types in this study, i.e. in gradient fractions XS - L and in non-enriched trophosome homogenate (Hom). ID count: number of identified proteins. Numbers are based on four biological replicates for sulfur-rich samples and three biological replicates for sulfur-depleted samples. Note that not all proteins were included in statistical analyses (StAn; see Methods for details). GF: gradient fractions.

| | sulfur-rich trophosome | | | | | sulfur-depleted trophosome | | | | | *total* |
|---|---|---|---|---|---|---|---|---|---|---|---|
| | Hom | XS | S | M | L | Hom | XS | S | M | L | |
| ID count | 1,151 | 1,022 | 1,296 | 1,603 | 1,722 | 1,017 | 1,099 | 1,260 | 1,605 | 1,572 | *1,946* |
| ID count (Hom only) | 1,151 | | | | | 1,017 | | | | | *1,223* |
| ID count (total all GF) | | 1,821 | | | | | 1,727 | | | | *1,898* |
| ID count (total all sample types) | 1,867 | | | | | 1,773 | | | | | *1,946* |
| proteins in StAn | | 940 | 1,081 | 1,135 | 1,134 | | 1,008 | 1,091 | 1,150 | 1,143 | *1,212* |
| proteins in StAn (total all GF) | | 1,135 | | | | | 1,151 | | | | *1,212* |

**Supplementary Table S7:** Total (summed up) relative abundance of *Endoriftia* proteins involved in specific metabolic categories in fractions XS, S, M and L in sulfur-rich (S-rich) *Riftia* specimens (average values, n=4) and sulfur-depleted (S-depl) *Riftia* specimens (average values, n=3). Only those 1,212 symbiont proteins presented in Supplementary Table S3a, which are included in the EdgeR statistical evaluation, are included (proteins with low abundance and/or only one or two replicate values were excluded). To allow comparison and summing of protein abundances across proteins within one sample, edgeR-RLE-corrected spectral count values were normalized a) to protein size, and b) to the sum of all proteins before summing up the proteins within categories (100% = all proteins in Supplementary Table S3a). These results indicate that morphological differences between individual symbiont differentiation stages are accompanied by a gradual change in metabolic function. During differentiation from small to large cells, *Riftia* symbionts rearrange their metabolic priorities, allocating resources to those processes that are most important in their respective life phase and role in the symbiosis.

| | S-rich | | | | S-depl | | | |
|---|---|---|---|---|---|---|---|---|
| **Category (based on KO, manually curated)** | **XS** | **S** | **M** | **L** | **XS** | **S** | **M** | **L** |
| Amino acid metabolism | 1.91 | 2.07 | 2.38 | 2.67 | 2.25 | 2.29 | 2.86 | 2.74 |
| Carbon metabolism | 16.55 | 17.52 | 17.84 | 18.49 | 17.61 | 17.71 | 18.30 | 18.21 |
| Cell cycle, cell division, cell shape | 0.47 | 0.53 | 0.66 | 0.71 | 0.57 | 0.58 | 0.68 | 0.70 |
| Cell wall | 1.55 | 1.67 | 1.90 | 1.91 | 1.83 | 1.88 | 2.07 | 1.97 |
| Chaperones, stress response | 3.69 | 3.40 | 3.03 | 3.15 | 3.88 | 3.98 | 3.41 | 3.30 |
| Cofactor and vitamin metabolism | 2.21 | 2.42 | 2.91 | 2.90 | 2.34 | 2.64 | 2.89 | 2.98 |
| DNA replication, recombination and repair | 0.97 | 0.86 | 1.11 | 1.02 | 1.11 | 1.07 | 1.28 | 1.28 |
| Energy metabolism | 6.31 | 6.90 | 7.77 | 7.65 | 6.45 | 6.95 | 7.21 | 7.46 |
| Genetic information processing | 2.51 | 2.56 | 2.89 | 2.81 | 2.69 | 2.78 | 3.01 | 2.97 |
| Lipid metabolism | 0.85 | 0.89 | 1.02 | 1.09 | 0.90 | 0.92 | 1.04 | 1.06 |
| Nitrogen metabolism | 3.46 | 3.51 | 3.86 | 3.63 | 3.29 | 3.64 | 3.68 | 4.01 |
| Nucleic acids metabolism | 1.55 | 1.61 | 1.78 | 1.91 | 1.78 | 1.72 | 1.90 | 1.95 |
| Other functions (including defense, secondary metabolism) | 0.66 | 0.67 | 0.74 | 0.74 | 0.69 | 0.72 | 0.84 | 0.86 |
| Protein folding and processing | 1.79 | 1.89 | 2.25 | 2.37 | 2.18 | 2.26 | 2.39 | 2.46 |
| Secretion, pilus, chemotaxis | 2.16 | 2.15 | 2.45 | 2.32 | 2.28 | 2.42 | 2.46 | 2.59 |
| Signaling | 0.61 | 0.68 | 1.03 | 1.04 | 0.82 | 0.88 | 1.23 | 1.29 |
| Sulfur metabolism | 21.73 | 23.03 | 21.83 | 21.81 | 18.54 | 19.08 | 18.42 | 18.60 |
| Transcription | 3.39 | 2.88 | 3.09 | 3.00 | 3.26 | 3.31 | 3.16 | 3.44 |
| Translation | 9.05 | 9.42 | 9.94 | 9.61 | 8.70 | 9.26 | 8.24 | 9.75 |
| Transporters | 13.86 | 10.91 | 6.87 | 6.42 | 13.66 | 10.83 | 9.80 | 7.28 |
| Unknown or general function prediction only | 4.69 | 4.41 | 4.65 | 4.74 | 5.16 | 5.08 | 5.11 | 5.12 |

**Supplementary Table S8**: *Riftia* trophosome homogenate and gradient fractions enriched in small and large symbionts, respectively, were stained with Syto9 and subjected to flow cytometry analysis in a FACSAria high-speed cell sorter with 488 nm excitation (see Methods for details). Two cell populations were identified, Pop1 and Pop2, which correspond to smaller and larger symbiont cells, respectively (see main text and Supplementary Figure S2). Median fluorescence intensity (FI) per particle, a measure of DNA content per cell, was compared between the two populations 1 and 2 to quantify differences in genome copy number between smaller and larger symbionts (column "ratio"). Note that FI ratios were not calculated for samples consisting of sorted populations (bottom rows), because these samples contained high cell numbers of either of the two populations, but very low cell numbers of the respective other population, preventing meaningful comparison. Analyses were performed with samples from two *Riftia* specimens (two biological replicates, BR).

| | | Population 1 | | | | Population 2 | | | | ratio |
|---|---|---|---|---|---|---|---|---|---|---|
| Sample description | Sample name | Count | Freq. of Parent | Mean FI | Median FI | Count | Freq. of Parent | Mean FI | Median FI | Median FI Pop2:Pop1 |
| Trophosome homogenate (BR 1) | 20200212_Riftia 19 HOM FA RNase_Syto9.fcs | 2398 | 12 | 1212 | 1097 | 2119 | 10.6 | 7341 | 7033 | 6.41 |
| Trophosome homogenate (BR 2) | 20200212_Riftia 21 HOM FA RNase_Syto9.fcs | 1802 | 9.01 | 253 | 186 | 1444 | 7.22 | 3401 | 3221 | 17.32 |
| Gradient fractions enriched in small symbionts (BR 1) | 20200212_Riftia 19 RZ07 FA RNase_Syto9.fcs | 4484 | 43.8 | 1373 | 1136 | 53 | 0.52 | 10317 | 9136 | 8.04 |
| | 20200212_Riftia 19 RZ08 FA RNase_Syto9.fcs | 9085 | 45.4 | 1395 | 1097 | 203 | 1.02 | 9828 | 8716 | 7.95 |
| | 20200212_Riftia 19 RZ09 FA RNase_Syto9.fcs | 8547 | 42.7 | 1470 | 1150 | 294 | 1.47 | 9260 | 8063 | 7.01 |
| Gradient fractions enriched in small symbionts (BR 2) | 20200212_Riftia 21 RZ07 FA RNase_Syto9.fcs | 2946 | 14 | 685 | 577 | 70 | 0.33 | 5748 | 4407 | 7.64 |
| | 20200212_Riftia 21 RZ08 FA RNase_Syto9.fcs | 2672 | 13.4 | 432 | 359 | 64 | 0.32 | 3832 | 2860 | 7.97 |
| | 20200212_Riftia 21 RZ09 FA RNase_Syto9.fcs | 2658 | 13.3 | 501 | 431 | 57 | 0.29 | 3466 | 2712 | 6.29 |
| Gradient fractions enriched in large symbionts (BR 1) | 20200212_Riftia 19 RZ19 FA RNase_Syto9.fcs | 2669 | 13.3 | 2549 | 1994 | 3003 | 15 | 11734 | 10459 | 5.25 |
| | 20200212_Riftia 19 RZ21 FA RNase_Syto9.fcs | 2910 | 14.5 | 2125 | 1686 | 3457 | 17.3 | 10249 | 9074 | 5.38 |
| | 20200212_Riftia 19 RZ22 FA RNase_Syto9.fcs | 2097 | 10.5 | 1957 | 1535 | 2669 | 13.3 | 10218 | 9534 | 6.21 |
| Gradient fractions enriched in large symbionts (BR 2) | 20200212_Riftia 21 RZ19 FA RNase_Syto9.fcs | 1242 | 6.21 | 735 | 451 | 2408 | 12 | 6746 | 5750 | 12.75 |
| | 20200212_Riftia 21 RZ20 FA RNase_Syto9.fcs | 1123 | 5.62 | 705 | 424 | 2026 | 10.1 | 7695 | 6792 | 16.02 |
| | 20200212_Riftia 21 RZ22 FA RNase_Syto9.fcs | 1633 | 8.16 | 893 | 497 | 2043 | 10.2 | 11365 | 10723 | 21.58 |
| Pop 1 sorted from trophosome homogenate | 20200212_small Syto sorted.fcs | | 49.9 | 485 | 419 | | 0.25 | 7737 | 7269 | |
| Pop 2 sorted from trophosome homogenate | 20200212_large Syto sorted.fcs | | 7.61 | 993 | 527 | | 49.6 | 3860 | 3708 | |

| | |
|---|---|
| **average ratio BR 1:** | **6.61** |
| standard deviation BR 1: | 1.04 |
| **average ratio BR 2:** | **12.79** |
| standard deviation BR 2 | 5.35 |
| **average ratio (total):** | **9.70** |
| standard deviation (total): | 4.94 |

**Supplementary Table S9:** Proteins identified as likely involved in dissimilatory sulfur metabolism in *Ca.* E. persephone after Blast-comparison against proteins identified in the literature: Weissgerber T, Sylvester M, Kröninger L, Dahl C. 2014. A comparative quantitative proteomic study identifies new proteins relevant for sulfur oxidation in the purple sulfur bacterium *Allochromatium vinosum*. Appl Environ Microbiol 80:2279–2292.; Rodriguez J, Hiras J, Hanson TE. 2011. Sulfite oxidation in *Chlorobaculum tepidum*. Front Microbiol 2:1–7.; Gregersen LH, Bryant DA, Frigaard NU. 2011. Mechanisms and evolution of oxidative sulfur metabolism in green sulfur bacteria. Front Microbiol 2:116. Significant - protein abundance significantly different between fractions containing symbionts of different size (see Methods for details on statistical analysis). Y - yes, N - no, M - may be.

| **Accession** | **Protein annotation** | **involved in sulfur oxidation?** | **detected as protein?** | **significant?** | **general abundance trend (if significant)** | |
|---|---|---|---|---|---|---|
| | | | | | **S-rich** | **S-depleted** |
| Sym_2601635419 | Adenylylsulfate reductase subunit alpha AprA | Y | Y | Y | small>large | |
| Sym_2601635420 | Adenylylsulfate reductase subunit beta AprB | Y | Y | N | | |
| Sym_EGV50053.1 | adenylylsulfate reductase, alpha subunit AprA | Y | Y | N | | |
| Sym_EGV52780.1 | anaerobic dimethyl sulfoxide reductase chain B/ SreB/SoeB/ PSRLC3 | M | N | N | | |
| Sym_2601636305 | DsrA | Y | Y | N | | |
| Sym_EGV52261.1 | DsrA | Y | Y | N | | |
| Sym_EGV52262.1 | DsrB | Y | Y | N | | |
| Sym_EGV52266.1 | DsrC | Y | Y | Y | small>large | small>large |
| Sym_EGW53956.1 | DsrC | Y | N | N | | |
| Sym_EGV52257.1 | DsrC family protein | Y | Y | Y | small>large | |
| Sym_EGV52361.1 | DsrC/ DsrC family | Y | Y | Y | | small>large |
| Sym_EGV50535.1 | DsrC/ DsrC-like | Y | Y | N | | |
| Sym_EGV52263.1 | DsrE | Y | Y | N | | |
| Sym_EGV50105.1 | DsrE2 | Y | N | N | | |
| Sym_EGV51796.1 | DsrE2 | Y | Y | Y | large>small | |
| Sym_EGV52264.1 | DsrF | Y | Y | Y | | small>large |
| Sym_EGV52265.1 | DsrH | Y | Y | N | | |
| Sym_EGV52270.1 | DsrJ | Y | N | N | | |
| Sym_EGV52268.1 | DsrK | Y | Y | Y | large>small | large>small |
| Sym_EGV52269.1 | DsrL | Y | Y | N | | |
| Sym_EGV52267.1 | DsrM | Y | Y | N | | |
| Sym_EGV52273.1 | DsrN | Y | N | N | | |
| Sym_EGW53659.1 | DsrN | Y | Y | N | | |
| Sym_2601636291 | DsrN / cobyrinate a,c-diamide synthase | M | Y | N | | |
| Sym_2601636293 | DsrO | Y | Y | N | | |
| Sym_EGV52272.1 | DsrP | Y | N | N | | |
| Sym_EGV52275.1 | DsrR | Y | Y | N | | |
| Sym_EGV52276.1 | DsrS | Y | Y | Y | large>small | |
| Sym_2601634706 | FccA | Y | N | N | | |
| Sym_EGV51006.1 | FccA | Y | Y | N | | |
| Sym_EGV52863.1 | FccA | Y | Y | N | | |
| Sym_EGV52186.1 | FccA (putative) | Y/M | N | N | | |
| Sym_EGV49859.1 | FccB | Y | N | N | | |
| Sym_EGV51007.1 | FccB | Y | Y | Y | small>large | |
| Sym_EGV50679.1 | PhsC; thiosulfate reductase cytochrome b subunit | M | Y | Y | small>large | small>large |
| Sym_EGV50955.1 | putative SoxL | Y/M | Y | Y | | large>small |
| Sym_EGV51355.1 | putative SoxW type thioredoxin | M | Y | N | | |
| Sym_2601633320 | QmoA | Y | Y | Y | large>small | |
| Sym_EGV50291.1 | QmoA | Y | Y | N | | |
| Sym_EGV52354.1 | QmoA | Y | Y | Y | large>small | |
| Sym_EGV52355.1 | QmoB | Y | Y | N | | |
| Sym_EGV52356.1 | QmoC | Y | Y | Y | | large>small |
| Sym_EGV50779.1 | rhodanese-like protein | M | Y | N | | |
| Sym_EGV49918.1 | Sgp protein | Y | Y | Y | small>large | small>large |
| Sym_EGV51298.1 | SgpA | Y | Y | N | | |
| Sym_EGV51608.1 | SgpB | Y | Y | N | | |
| Sym_EGW54364.1 | SgpB | Y | Y | Y | small>large | small>large |
| Sym_EGV50398.1 | SgpC | Y | N | N | | |
| Sym_EGW54499.1 | SgpC | Y | N | N | | |
| Sym_EGV51976.1 | SoeA | Y | Y | Y | | large>small |
| Sym_EGV51975.1 | SoeB | Y | Y | N | | |
| Sym_EGV51974.1 | SoeC | Y | Y | N | | |
| Sym_EGV50245.1 | SoeC/ anaerobic dimethyl sulfoxide reductase, A subunit, DmsA/YnfE family | M | N | N | | |
| Sym_EGV50426.1 | SoxA | Y | Y | N | | |
| Sym_2601635312 | SoxB | Y | Y | N | | |
| Sym_EGV50931.1 | SoxB | Y | Y | N | | |
| Sym_EGV50425.1 | SoxK | Y | Y | N | | |
| Sym_EGV50424.1 | SoxL | Y | Y | N | | |
| Sym_EGV52219.1 | SoxL/ rhodanese domain protein | M | Y | N | | |
| Sym_EGV50427.1 | SoxX | Y | Y | N | | |
| Sym_EGV52247.1 | SoxY | Y | Y | N | | |
| Sym_EGV52246.1 | SoxZ | Y | Y | Y | large>small | |
| Sym_EGV51808.1 | SqrF | Y | Y | N | | |
| Sym_EGV51162.1 | SreA | Y | Y | N | | |
| Sym_2601636275 | sreA; sulfur reductase molybdopterin subunit | Y | N | N | | |
| Sym_EGV51161.1 | SreB | Y | N | N | | |
| Sym_EGV51160.1 | SreC | Y | N | N | | |
| Sym_EGV50710.1 | Sulfate adenylyltransferase Sat | Y | Y | N | | |
| Sym_EGV49794.1 | Sulfate permease | Y | Y | N | | |
| Sym_2601634801 | sulfate permease (SulP) | Y | Y | Y | large>small | large>small |
| Sym_EGV52704.1 | Sulfate transporter | Y | Y | N | | |
| Sym_EGV50940.1 | Sulfate transporter cysZ | Y | Y | N | | |
| Sym_EGV50288.1 | Sulfhydrogenase 1 subunit beta hydB | M | Y | N | | |
| Sym_EGV50287.1 | Sulfhydrogenase 1 subunit gamma hydG/ Anaerobic sulfite reductase subunit B ArsB | M | Y | Y | | large>small |
| Sym_EGV50140.1 | Sulfide-quinone reductase SqrD | Y | N | N | | |
| Sym_2601635970 | Sulfur carrier protein DsrE2 | Y | N | N | | |
| Sym_EGV51798.1 | Sulfurtransferase Alvin_2599 (Rhd_2599) | Y | Y | N | | |
| Sym_EGV52654.1 | thiosulfate sulfurtransferase | M/N | Y | N | | |
| Sym_EGV50246.1 | TtrB; tetrathionate reductase subunit B/ SoeB/ SreB | M | N | N | | |
| Sym_EGV51797.1 | TusA | Y | Y | Y | small>large | small>large |
