## Supplementary figures and images for "Metabolic differences between symbiont subpopulations in the deep-sea tubeworm *Riftia pachyptila*"

### Supplementary Figure S5

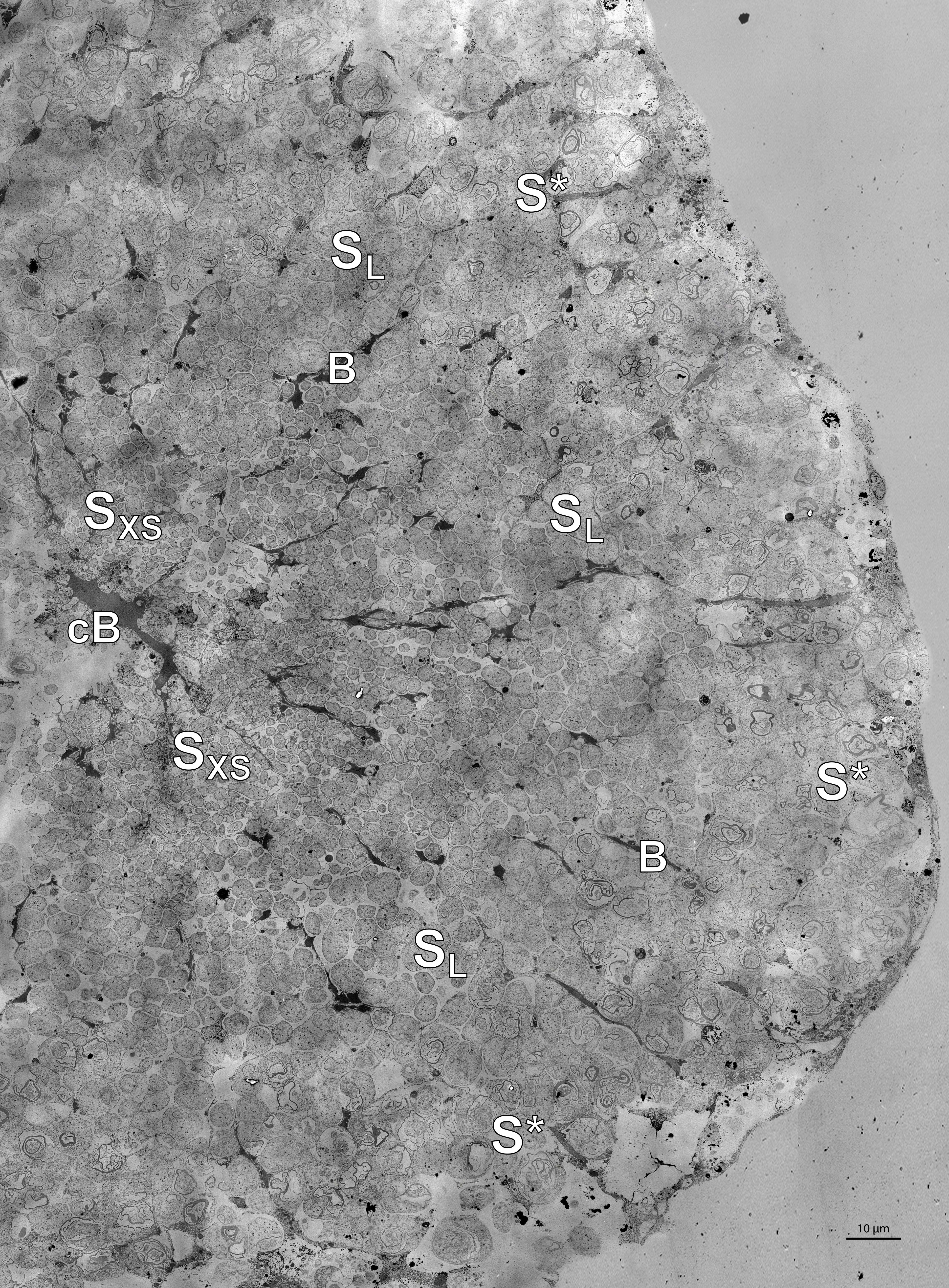
